# Discovery and characterization of a copper-binding carbohydrate-binding module (CBM) regulating the activity of lytic polysaccharide monooxygenases

**DOI:** 10.1101/2025.06.11.659075

**Authors:** Zarah Forsberg, Anton A. Stepnov, Ole Golten, Esteban Lopez-Tavera, Iván Ayuso-Fernández, Vincent G. H. Eijsink

**Author notes:** To whom correspondence should be addressed: Zarah Forsberg & Vincent G.H. Eijsink, **Email:** &.

## Abstract

Lytic polysaccharide monooxygenases (LPMOs) are monocopper enzymes that hydroxylate recalcitrant polysaccharides such as cellulose. Like other redox enzymes, LPMOs face challenges in handling the reactive oxygen species generated at their active site, which must be controlled to prevent off-pathway reactions that lead to enzyme inactivation. In the case of LPMOs, oxidative damage may be self-reinforcing because free copper released from damaged catalytic centers will promote abiotic redox reactions that generate reactive oxygen species. Here we show that some members of a widely spread family of carbohydrate-binding modules (CBM2s) have evolved the ability to bind copper and that this ability is exclusively found in CBM2s that are appended to LPMOs. We show that the copper site in these CBM2s protects the LPMO from inactivation both by scavenging free copper, preferably Cu(I), and by interacting directly with the reduced catalytic copper site of the LPMO, thus preventing the enzyme from engaging in off-pathway reactions. These effects are demonstrated by studies on the redox stability of a series of engineered LPMO variants as well as AlphaFold3 models of the CBM2-containing enzymes. Interestingly, the copper site and the cellulose-binding surface are located on different sides of these CBM2s, enabling a mode of action in which the CBM inhibits potentially damaging LPMO activity in the absence of substrate, while such inhibition would be relieved upon binding of the CBM2 to cellulose. These findings show that CBMs have biologically relevant functions beyond carbohydrate-binding and reveal a mechanism for substrate-dependent regulation of LPMO reactivity.

## Introduction

Copper is an essential micronutrient for bacteria, acting as a cofactor for redox-active cuproenzymes. However, the redox properties that make copper valuable in these metalloproteins also cause oxidative damage. The reaction of Cu(I) with hydrogen peroxide, followed by the re-reduction of Cu(II) by superoxide (a process similar to Fenton and Haber-Weiss reactions catalyzed by iron), produces hydroxyl radicals that can damage proteins, lipids, and nucleic acids [1, 2]. To harness the beneficial roles of copper while mitigating its potential toxicity, organisms have evolved various mechanisms for copper homeostasis, ensuring a balance that supports essential metabolic processes without causing oxidative damage [3-8].

Lytic polysaccharide monooxygenases (LPMOs) are oxidoreductases that utilize a single copper ion as a cofactor [9] to oxidatively cleave recalcitrant polysaccharides such as cellulose and chitin [10-12, 9], making these substrates more accessible to the enzymatic depolymerization that is needed to generate compounds, such as glucose, that feed into metabolic pathways [13]. LPMOs are currently classified into eight families within the Carbohydrate-Active enZymes database (CAZy; http://www.cazy.org/; [14]), namely auxiliary activity (AA) families 9–11 and 13–17. These enzymes are predominantly found in fungi and bacteria but are also present in insects [15], plants [16], and viruses [17]. While LPMOs have mainly been studied in the context of biomass degradation, recent research has revealed a broader range of potential LPMO functions [18] including roles in pathogenesis [19-24], commensalism [25, 26] and cell wall remodeling [27, 28].

Despite their name, LPMOs are primarily peroxygenases that utilize hydrogen peroxide (H_2_O_2_) as a cosubstrate (H_2_O_2_ + R-H → R-OH + H_2_O) [29], contrary to the earlier assumption that molecular oxygen (O_2_) played this role [10]. Depending on reactant levels and enzyme-specific redox properties, LPMOs can also exhibit reductant oxidase (O_2_ + RedH_2_ → H_2_O_2_ + Red) and peroxidase (H_2_O_2_ + RedH_2_→ 2 H_2_O + Red) activities. The latter two pathways are more prominent in the absence of a polysaccharide substrate. The productive peroxygenase reaction involves reductive activation of the enzyme [Cu(II) + e^-^ → Cu(I)], and homolytic cleavage of H_2_O_2_ that eventually leads to the formation of a copper-oxyl intermediate [29-33]. Like other oxidoreductases, LPMOs are prone to oxidative damage [29, 34-36]. Excess H_2_O_2_ or the absence of substrate can drive autocatalytic oxidation of the Cu-coordinating histidines via the peroxidase pathway, leading to enzyme inactivation [29]. Once the histidines are damaged, copper leaks from the active site [37, 36], which may accelerate the damaging peroxidase reaction as unbound copper rapidly reacts with O_2_ and reductants to generate high levels of H_2_O_2_. This phenomenon has been described as a self-reinforced inactivation process [37].

Nonetheless, LPMOs have evolved mechanisms to resist damage and prevent inactivation. Fungal LPMOs feature methylation of the N-terminal histidine, enhancing H_2_O_2_ tolerance and stability [9, 38, 34]. Additionally, in both bacterial and fungal LPMOs, chains of tyrosine and tryptophan residues provide “hole-hopping” pathways that divert radicals away from the active site [39-41]. Comparative studies have suggested that fungal LPMOs may on average achieve some 100 peroxidase turnovers before inactivation, while this number is lower (10 – 40) for bacterial LPMOs [34].

Carbohydrate-binding modules (CBMs) have long been known for enhancing polysaccharide conversion by glycoside hydrolases [42-45], and recent findings suggest this also applies to LPMOs [46-50]. Importantly, LPMOs linked to CBMs are generally more stable than single-domain LPMOs as being close to the substrate promotes productive consumption of H_2_O_2_ over potentially damaging futile peroxidase reactions [29, 49, 50]. Inspired by previously unexplained effects of CBM removal on LPMO activity and stability ([37]; see below) and the advent of AlphaFold 3 [51], we have discovered that certain CBMs, exclusively tethered to LPMOs, contain copper-binding sites. We have investigated the occurrence and binding properties of these copper sites as well as their impact on LPMO performance. We have studied two seemingly similar CBM-containing LPMOs, one with the newly discovered copper site and one without it, and used CBM swapping as well as site-directed mutagenesis to introduce and eliminate copper-binding capacity. Our findings reveal that these copper-binding CBMs help protect the LPMO from damaging off-pathway reactions, offering new perspectives on the roles of CBMs and on the determinants of LPMO performance.

## Results & Discussion

### Discovery of an additional copper binding site in *Sc*LPMO10C

The cellulose-active LPMO from *Streptomyces coelicolor* A3(2), called *Sc*LPMO10C, is one of the first and most extensively studied LPMOs to date [11, 52, 29, 46, 53, 37, 54, 55]. It is strictly C1-oxidizing and its catalytic domain is attached, via a flexible linker of approximately 30 amino acids [54], to a CBM from family 2 (hereinafter referred to as *Sc*CBM2), known to bind cellulose [46]. In a previous study, we observed that the apparent oxidase activity of the catalytic domain of *Sc*LPMO10C, hereafter referred to as *Sc*AA10 or CD, is three-fold higher than that of the full-length enzyme [37]. Furthermore, in follow-up studies, we noted that, in reactions without substrate, the CD showed typical rapid self-reinforced inactivation, which involves release of copper from damaged enzymes, whereas the full-length enzyme did not. These remarkable effects of the CBM in the absence of substrate led us to consider whether the CBM could have affinity for copper and/or whether the CBM could interact with the catalytic copper site to modulate its reactivity. Interactions between the CD and the CBM have not been observed using NMR [46, 54] or SAXS [54], but the hypothesis remains plausible given the length of the linker and given a possible dependence on the redox state of the LPMO. Notably, the NMR and SAXS studies were conducted with the *apo* form of the enzyme in the absence of reductants.

Strikingly, structure predictions with AlphaFold3 revealed that the two domains interact, but only in the presence of copper (Fig. 1). Although still relatively new, AlphaFold3 has demonstrated promising accuracy in modelling metal coordination, which supports the plausibility of the predicted copper-dependent interaction [51]. The AlphaFold structure shows a metal-binding site composed of the histidine brace in the catalytic domain and, primarily, His288 in the CBM. His288 is part of a structural motif also including two methionines (Met266 and Met268), resembling the Cu(I)-binding sites in the periplasmic copper-trafficking proteins CusF and CopC [56, 57]. Studies with recombinantly produced *Sc*CBM2, which are discussed in detail below, showed that this CBM2, most remarkably, indeed binds copper.

**Figure 1.**
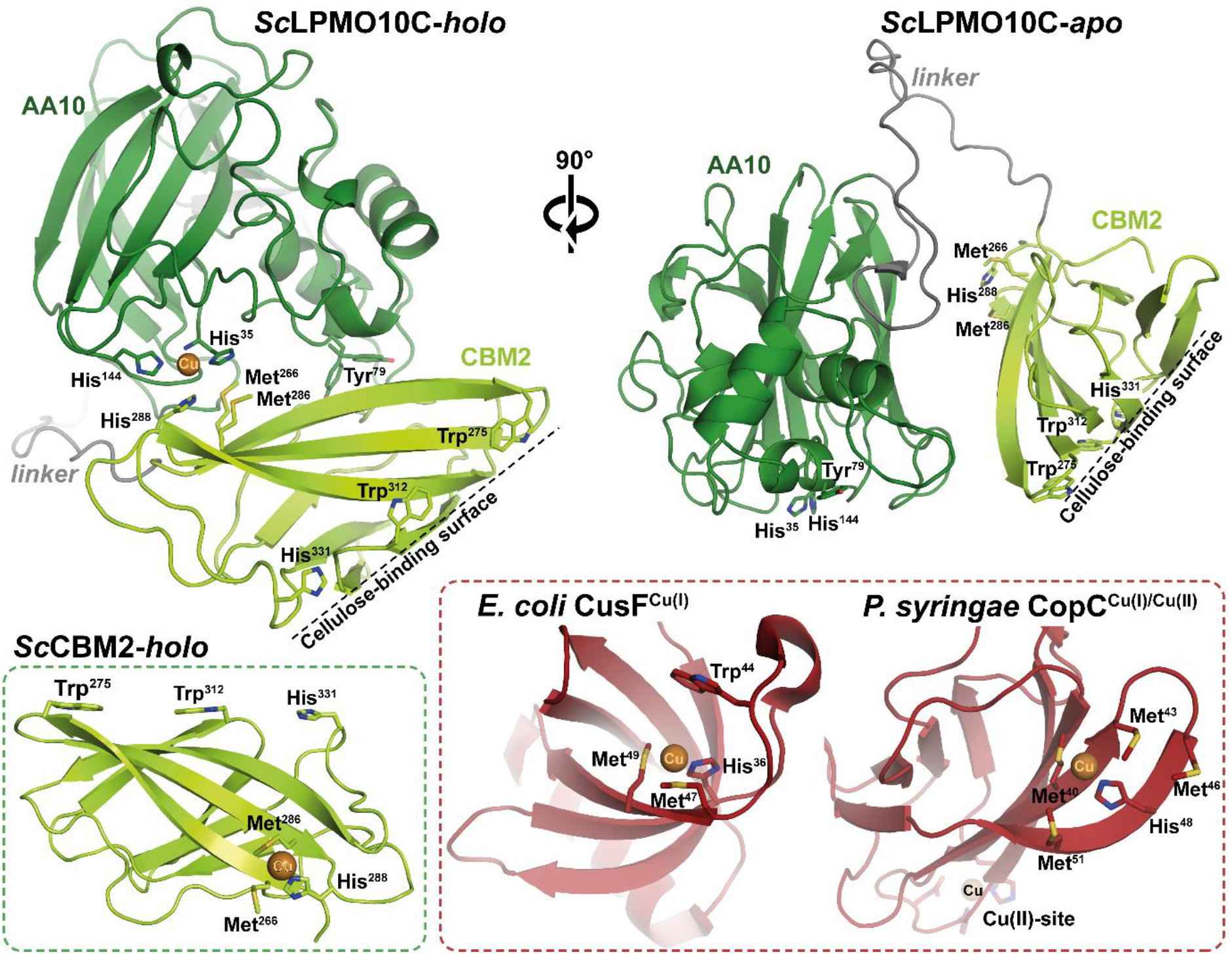
Predicted structure of *Sc*LPMO10C generated by AlphaFold3 in the presence (*holo*) and absence (*apo*) of copper. The structure on the left is Cu(II)-loaded *Sc*LPMO10C, highlighting an intramolecular interaction between the histidine brace (His35 and His144) of the AA10 and a potential copper-binding site on the CBM2 (Met266, Met286, and His288). This interaction is absent in the predicted structure of the *apo* protein, which is shown to the right, rotated 90° along the y-axis relative to the catalytic AA10 domain to make the CBM visible. The identified copper-binding site on the CBM2 (lower left structure) is similar to those found in periplasmic copper-trafficking proteins, where Cu(I) is coordinated by two methionines and one histidine. Examples include CusF (PDB: 2VB2) from *Escherichia coli* [58] and CopC (PDB: 2C9Q) from *Pseudomonas syringae pv. tomato* [59].

### A conserved MMH-motif occurs exclusively in CBM2s tethered to C1-oxidizing LPMOs acting on cellulose

In Nature, CBM2 domains are associated with a diverse range of bacterial CAZymes, such as various cellulases. An analysis of all CBM2-containing sequences available in InterPro (IPR001919; >26,000 sequences) was performed to identify the sequence motif MX_n_MXH (where X represents any amino acid). Structural verification using AlphaFold-predicted models revealed that the MMH motif, first identified in *Sc*CBM2, is exclusively found in CBM2s associated with AA10 domains (649 proteins), as well as in 34 putative proteins containing only a CBM2 and no additional domains (Fig. 2A). Of the 649 MMH-containing AA10-CBM2 proteins, 644 possessed a canonical arrangement of Arg, Glu, and Phe (REF motif) in the second coordination sphere of the copper in the AA10 domain [60, 61], suggesting that these all are *bonafide* C1-oxidizing cellulose-active LPMOs.

**Figure 2.**
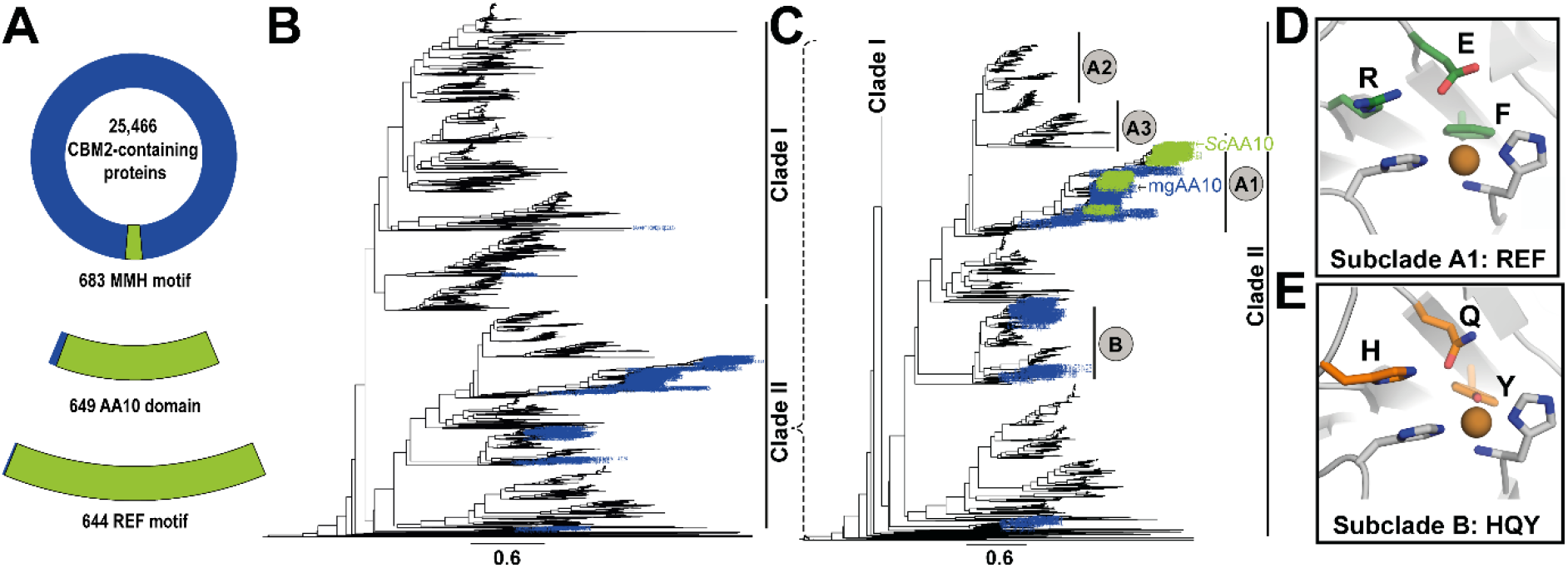
Co-occurrence of the MMH-motif in CBM2s and the second sphere REF-motif in C1-oxidizing cellulose-active AA10s. Panel A shows a pie chart illustrating all CBM2 domains identified in InterPro (IPR001919) and the occurrence of the MMH motif (∼2.7%). Among these MMH-containing CBM2s, 95% are associated with an AA10 domain, while the remaining 5% lack a catalytic domain. Of the AA10s associated with MMH-containing CBM2s, 99% contain the REF motif. Panel B shows a phylogenetic tree of AA10 LPMOs, generated using a dataset curated with the in-house script dbcan_curation.sh, based on sequences available in dbCAN (n=5433; catalytic domains only). Two major clades were identified: Clade I, containing chitin-active LPMOs, and Clade II (expanded in panel C), comprising LPMOs active on cellulose, chitin, or both. Sequences were aligned using MAFFT (L-INS-i option), and the tree was constructed with FastTree using default parameters. Panel C shows the subclades in Clade II: blue-labeled branches represent AA10-sequences linked to a CBM2, while green-labeled branches represent LPMOs with CBM2s containing the MMH motif. The MMH motif is only found in subclade A1, which harbors AA10 domains with the second sphere REF-motif (panel D), as opposed to the HQY-motif (panel E) that is observed in subclade B.

Among the entire pool of 5433 AA10 sequences available in dbCAN3 (assessed June 2023), 9% are linked to CBM2s (Fig. 2B). These CBM2s are predominantly associated with LPMOs from clade II, which contains all known cellulose-active AA10s [62, 63]. The cellulose-active enzymes in clade II (Fig. 2C) are separated into strict C1-oxidizing LPMOs (subclade A1) and LPMOs with mixed C1/C4 activity on cellulose in addition to C1-activity on chitin (subclade B). Strict C1-oxidizing LPMOs possess a second-sphere REF-motif (Arg-Glu-Phe; Fig. 2D), whereas C1/C4-oxidizing LPMOs feature the HQY-motif (His-Gln-Tyr; Fig. 2E). Analysis of all (130) AA10s included in dbCAN3 with an MMH-containing CBM2 showed that these CBM2s are exclusively associated with LPMOs containing the REF second sphere motif in subclade A1 (Fig. 2C). Approximately 81% of the LPMOs in subclade A1 (321 in total) contain a CBM2 and ∼40% of these CBM2s (130 in total) contain the MMH motif.

To further investigate the interaction of the MMH-containing CBM2 domains with their respective catalytic domains, we generated AlphaFold2 [64] models for all 130 sequences, as well as 28 models of AA10s with CBM2s lacking the MMH motif. The models of AA10s with MMH-containing CBMs consistently predicted an intramolecular interaction similar to that observed for *Sc*LPMO10C. In contrast, the models for AA10s with CBMs lacking the MMH motif displayed greater variability in terms of how the CBM2 is positioned relative to the catalytic domain (Fig. S1).

### Engineering the MMH-motif in two similar LPMOs with distinct CBM2s

To investigate the copper-binding function of the MMH-motif, the three residues in *Sc*LPMO10C were mutated to alanines (M266A/M286A/H288A) in both the full-length enzyme and the isolated CBM2 domain. In a complementary approach, the MMH motif was introduced into an LPMO that naturally lacks this motif. We chose mgLPMO10 for this purpose due to its phylogenetic proximity to *Sc*LPMO10C (Fig. 2) and similar C1-oxidizing activity [65]. The overall sequence identity between the two full-length enzymes is 56%, with 62% identity in the catalytic domains and 47% in the CBM2s. mgLPMO10 lacks the MMH residues, which are replaced by ART (Ala268-Arg288-Thr290). Site-directed mutagenesis was used to introduce the MMH-motif in both the full-length enzyme and the isolated CBM2 domain (mgCBM2). Additionally, the entire CBM2 domain was substituted between the two LPMOs to assess whether other residues beyond the MMH-motif contribute to the predicted interaction between the two domains. Next to this LPMO pair, we studied another novel LPMO with an expanded MMH motif (MMHH), called *Af*LPMO10B. Table 1 shows an overview of all the protein variants included in this study.

**Table 1.** Overview of protein variants produced and characterized in this study. Motifs in brackets, i.e., MMH and ART, represent wildtype motifs, whereas motifs without brackets are the result of one or more site-directed mutations in the CBMs. Asterisks indicate the number of mutations introduced.

| Protein | Domain | Mutation(s) |
| --- | --- | --- |
| ScLPMO10C<sup>(MMH)</sup> | AA10-CBM2 | - |
| ScAA10 | AA10 | - |
| ScCBM2<sup>(MMH)</sup> | CBM2 | - |
| ScAA10-mgCBM2<sup>(ART)</sup> | AA10-CBM2 | CBM substitution<br>(residues 263-364 replaced<br>by residues 265-363) |
| ScLPMO10C<sup>AMH</sup> | AA10-CBM2* | M266A |
| ScLPMO10C<sup>MMA</sup> | AA10-CBM2* | H288A |
| ScLPMO10C<sup>AAA</sup> | AA10-CBM2*** | M266A<br>M286A<br>H288A |
| ScCBM2<sup>AAA</sup> | CBM2*** | M266A<br>M286A<br>H288A |
| mgLPMO10<sup>(ART)</sup> | AA10-CBM2 | - |
| mgAA10 | AA10 | - |
| mgCBM2<sup>(ART)</sup> | CBM2 | - |
| mgAA10-ScCBM2<sup>(MMH)</sup> | AA10-CBM2 | CBM substitution<br>(residues 265-363 replaced<br>by residues 263-364) |
| mgLPMO10<sup>MMH</sup> | AA10-CBM2*** | A268M<br>R288M<br>T290H |
| mgCBM2<sup>MMH</sup> | CBM2*** | A268M<br>R288M<br>T290H |
| AfLPMO10B<sup>(MMHH)</sup> | AA10-CBM2 | - |
| AfCBM2<sup>(MMHH)</sup> | CBM2 | - |

Assessment of potential domain interactions between *Sc*AA10 (Fig. S2), mgAA10 (Fig. S3), and CBM variants containing AAA, ART, or MMH motifs using AlphaFold3 revealed that the MMH motif is essential for the interaction with the catalytic copper site. For example, upon insertion of the MMH motif in the CBM2 of mgLPMO10, this interaction was predicted, in contrast to the wildtype enzyme containing ART. Unexpectedly, while mutation of MMH in the CBM2 of *Sc*LPMO10C to AAA significantly reduced the likelihood of the interaction with the catalytic domain, it was still predicted, but only if the linker between the two domains was retained (Fig. S2). This suggests that other residues may contribute to this interaction. Of note, comparison of the predicted structures of the *Sc*AA10 and *Sc*CBM2^(MMH)^ domains with the crystal structure and the NMR structure of these domains yielded RMSD values of 0.23 Å and 1.57 Å, respectively, underpinning the reliability of the predictions (Fig. S4). It is worth noting that all the models of MMH-containing CBM2s binding to the catalytic copper site consistently show that only the histidine of the MMH motif interacts with the catalytic copper.

### CBM2s with the MMH motif bind copper with a preference for Cu(I)

Binding assays showed that the two wildtype CBM2s, *Sc*CBM2^(MMH)^, and mgCBM2^(ART)^, have similar K_d_ values for binding to Avicel, the cellulosic substrate used in all subsequent experiments (3.7 µM and 1.9 µM, respectively; Fig. S5). This is not surprising, since the cellulose-binding surfaces of the two CBM2s are very similar (Fig. S5), and show that the effects of the mutations discussed below, which do not affect the cellulose-binding surface, do not relate to changes in cellulose binding.

The presence of free copper promotes LPMO activity in reactions driven by the commonly used reductant ascorbic acid since free copper boosts abiotic oxidation of the reductant to generate the H_2_O_2_ that drives the LPMO reaction [66]. Copper binding by free *Sc*CBM2^(MMH)^ was initially suggested by the observation that addition of free *Sc*CBM2^(MMH)^ to an ascorbic acid-driven LPMO reaction abolishes the boosting effect of free copper (Fig. S6).

Copper binding by the four CBM variants studied here [*Sc*CBM2^(MMH)^, *Sc*CBM2^AAA^, mgCBM2^(ART)^, mgCBM2^MMH^] was assessed using bathocuproine disulfonate (BCS) as a Cu-probe [67, 68], which allows measuring binding of both Cu(I) and Cu(II) to proteins. The results (Fig. 3) demonstrate that *Sc*CBM2^(MMH)^ binds copper, with a preference for Cu(I), whereas wildtype mgCBM2^(ART)^ does not bind to either Cu(I) or Cu(II). Mutation of the MMH motif in *Sc*CBM2 to AAA resulted in loss of copper binding, while the reciprocal mutation in mgCBM2 (i.e., ART to MMH) led to apparent binding of both Cu(I) and Cu(II). While unraveling the true binding preferences of engineered, MMH-containing mgCBM2 requires additional validation (using anaerobic conditions), the results obtained with (wildtype) *Sc*CBM2^(MMH)^ are crystal clear: this CBM2 binds copper through the MMH site and prefers Cu(I).

**Figure 3.**
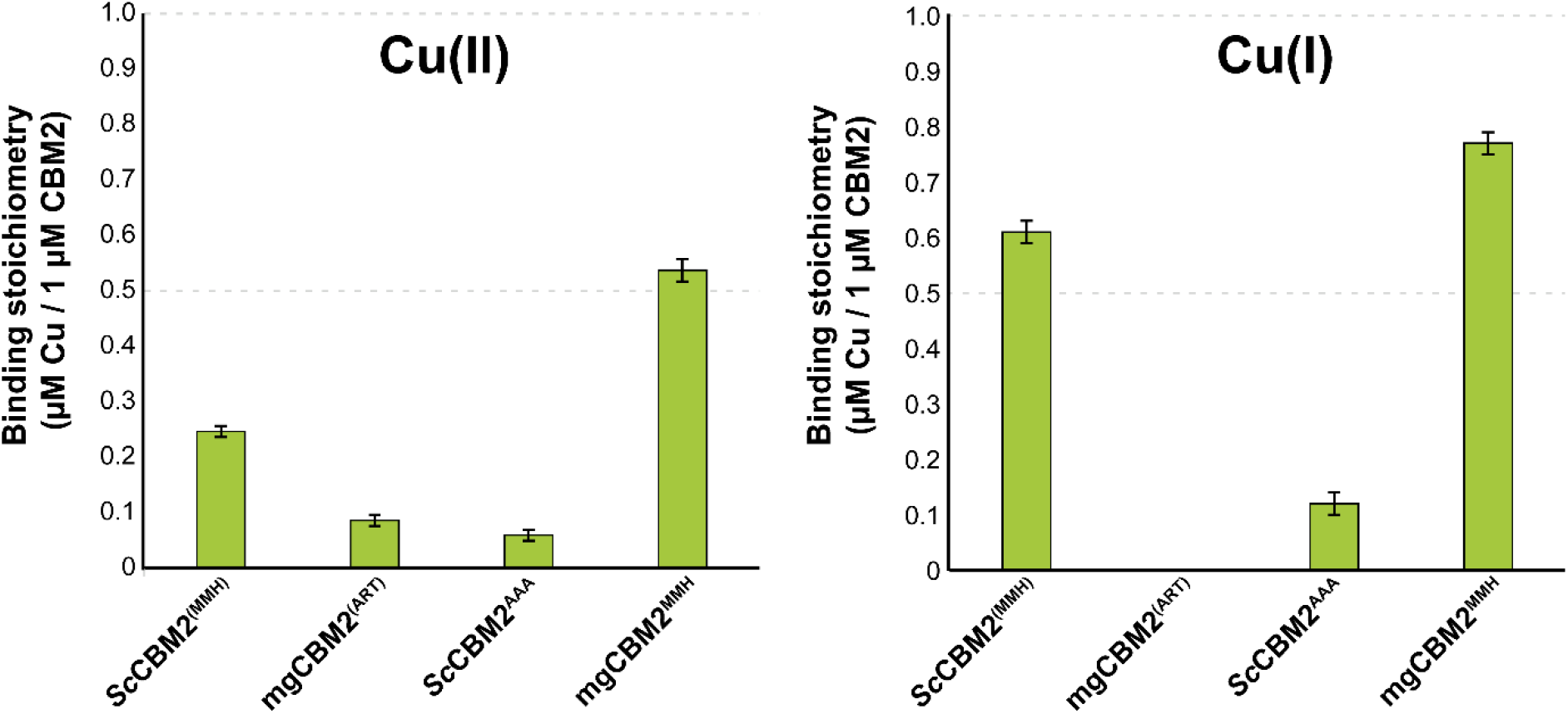
Binding of Cu(II) and Cu(I) by wildtype and mutant CBM2s. Copper binding was assessed under oxidizing (left) or reducing (right) conditions for the two wildtype CBM2s (*Sc*CBM2^(^MMH^)^ and mgCBM2^(^ART^)^) and the two mutated CBM2s (*Sc*CBM2AAA and mgCBM2MMH). For each assay, 4 µM CBM was incubated with 8 µM Cu(II)SO_4_, with or without 20 µM ascorbate, for 10 min in 50 mM sodium phosphate buffer (pH 6.0). After incubation, unbound copper was separated from the protein by ultrafiltration using a 3 kDa cut-off filter, and the supernatant containing the unbound copper was mixed with BCS and 40 µM ascorbate to assure full reduction and binding to BCS. Fluorescence (Ex/Em = 290/325 nm) was measured and converted to copper concentration using a standard curve [0-8 µM Cu(II)SO_4_] which was treated identically to the CBM samples. The y-axis represents the amount of bound Cu(II) or Cu(I), normalized to 1 µM of CBM2, indicating the binding stoichiometry for each condition. Error bars represent standard deviations (n = 3).

### Evaluating the effect of copper-binding CBMs on LPMO performance – cellulose degradation

Before conducting experiments, all LPMOs were loaded with copper [Cu(II)] as described in Materials and Methods. To confirm that *Sc*LPMO10C and mgLPMO10 variant preparations were free of excess copper and that their CBMs did not contain copper, ICP-MS analysis was performed (Table S1). The analysis revealed that all LPMOs contained sub-stoichiometric amounts of copper, ranging from 0.6 to 0.9 µM copper per 1 µM LPMO. The free CBM2s showed negligible levels of bound copper (Table S1).

LPMO activity on Avicel was assessed in reductant-driven reactions (also known as “H_2_O_2_-limiting” or “apparent monooxygenase” conditions). In these reactions, H_2_O_2_ is generated by the oxidase activity of the LPMO and by abiotic oxidation of the reductant, the latter typically catalyzed by trace metal ions at physiological pH. Fig. 4 shows the clear and well known [46, 48-50] advantage of having a substrate-binding CBM. The CBM promotes the use of available H_2_O_2_ in productive reactions with substrate, rather than in damaging peroxidase reactions, which enhances enzyme stability and leads to high final product levels. This type of progress curve is obtained regardless of the presence of the MMH-motif and only when the CBM is covalently bound to the CD. Reactions with the truncated enzyme, *Sc*AA10, show very different progress curves, which has been observed and explained previously [37]. This CD alone binds weakly to the substrate, which leads to enhanced oxidase activity [69] while increasing the chance that the generated H_2_O_2_ is consumed in the peroxidase reaction. As a result, the reaction with *Sc*AA10 is fast, but the enzyme inactivates quickly, and the final product level is low (Fig. 4).

**Figure 4.**
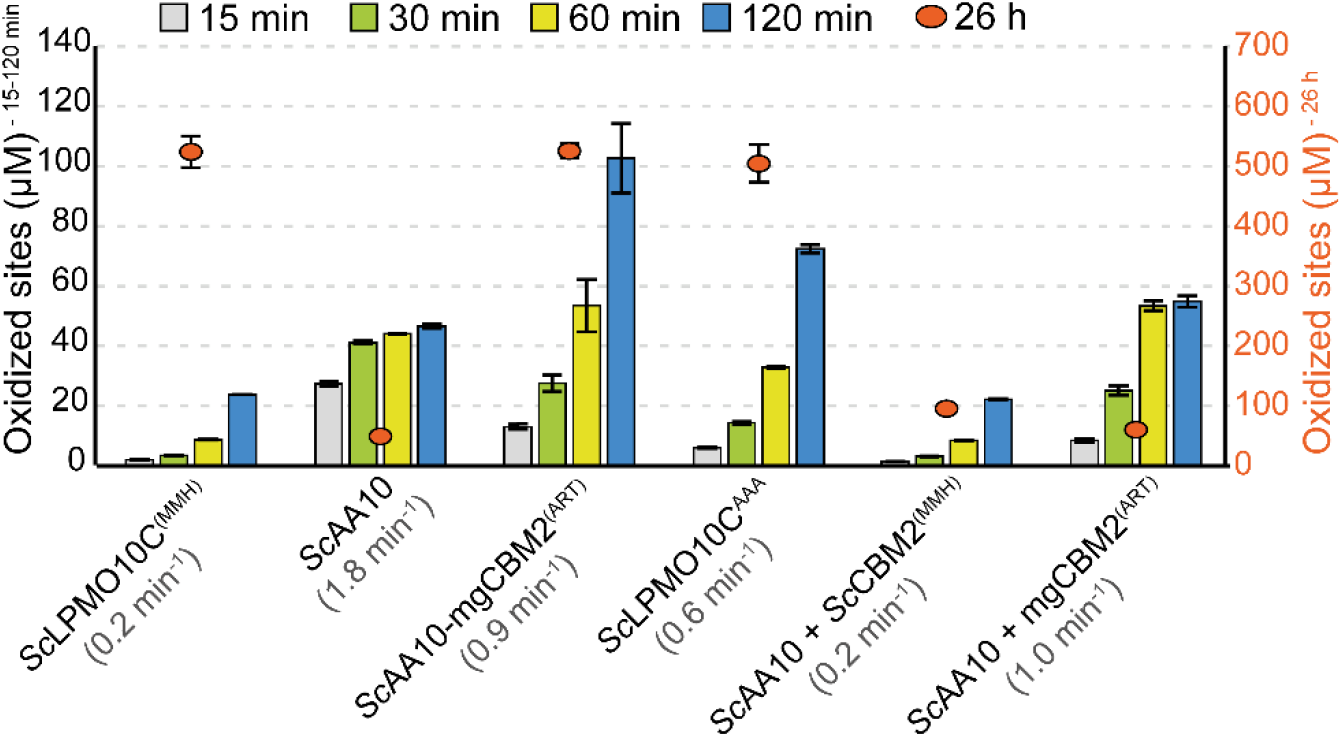
Quantification of soluble products from Avicel degradation by *Sc*LPMO10C variants and domain combinations over time. Reactions containing 1 µM LPMO were incubated with 10 g/L Avicel at 40°C in 50 mM sodium phosphate buffer (pH 6.0), with 1 mM ascorbic acid, for up to 26 hours. At various time points, samples were taken, and reactions were stopped by vacuum filtration. The soluble oxidized products were then converted to oxidized dimers and trimers, by the *Thermobifida fusca* endoglucanase Cel6A (*Tf*Cel6A), which were quantified to yield oxidized sites (left y-axis for the early time points; right y-axis for the 26-h time point). The initial rates (shown in brackets for each reaction) were estimated from the linear phase of the reaction, which varied significantly between enzymes and enzyme combinations. Error bars represent standard deviations (n = 3). Similar data for mgLPMO10, showing similar trends, are shown in Fig. S7.

Importantly, Fig. 4 shows two features that relate to the copper-binding ability of the CBM2. Firstly, addition of *Sc*CBM2^(MMH)^ drastically slows down Avicel oxidation by *Sc*AA10 and leads to slower inactivation and an increase in the final product level, whereas this effect is much less pronounced when adding mgCBM2^(ART)^. This shows that, in reactions with added *Sc*CBM2^(MMH)^, binding of the free copper that is released by damaged LPMOs will reduce the copper-catalyzed increase in H_2_O_2_ production, which will reduce the reaction rate and dampen the self-reinforcing inactivation process. Secondly, Fig. 4 shows that altering the MMH copper site in *Sc*LPMO10C to ART (domain swapping; *Sc*AA10-mgCBM2^(ART)^) or AAA (mutagenesis; *Sc*LPMO10C^AAA^) leads to clearly higher reaction rates. This may be due to less scavenging of free copper in the two mutants, but also suggests that the MMH-containing CBM2 domain indeed interacts with the catalytic copper site, which would reduce enzyme activity. In this respect, it is worth noting that AlphaFold3 predicts that the CBM still has some affinity for the CD in *Sc*LPMO10C^AAA^ (Fig. S2), whose progress curve lies in between those for the wild-type enzyme (slower; interaction predicted) and *Sc*AA10-mgCBM2^(ART)^) (faster; interaction not predicted). Notably, similar effects were observed for the corresponding mgLPMO10 variants, as shown in Fig. S7.

### Evaluating the effect of copper-binding CBMs on LPMO performance – copper reactivity

It is conceivable that binding of the MMH motif to the LPMO copper site reduces the redox activity of that copper site. This was first assessed by examining whether the MMH-motif influences the oxidase activity of the LPMO. Of note, this reaction requires multiple steps involving both the Cu(I) and the Cu(II) state and the rate-limiting step remains uncertain [33, 70, 71]. The removal of the MMH-motif in *Sc*LPMO10C, either through deletion (*Sc*AA10), point mutations (*Sc*LPMO10C^AAA^) or complete CBM2 substitution (*Sc*AA10-mgCBM2^(ART)^), resulted in slightly higher oxidase activity compared to the wildtype enzyme (Fig. 5), which could be taken to confirm that the two domains indeed interact, limiting the reaction, but could also be due to reduced binding of free copper ions in the reaction. Importantly, an opposite effect was not observed upon introducing the MMH motif into mgLPMO10 (as in mgAA10-*Sc*CBM2^(MMH)^ and mgLPMO10^MMH^ (Fig. S8). All in all, the effects were small, suggesting that the two domains do not interact when the LPMO is in the state involved in the rate-limiting step of the oxidase reaction.

**Figure 5.**
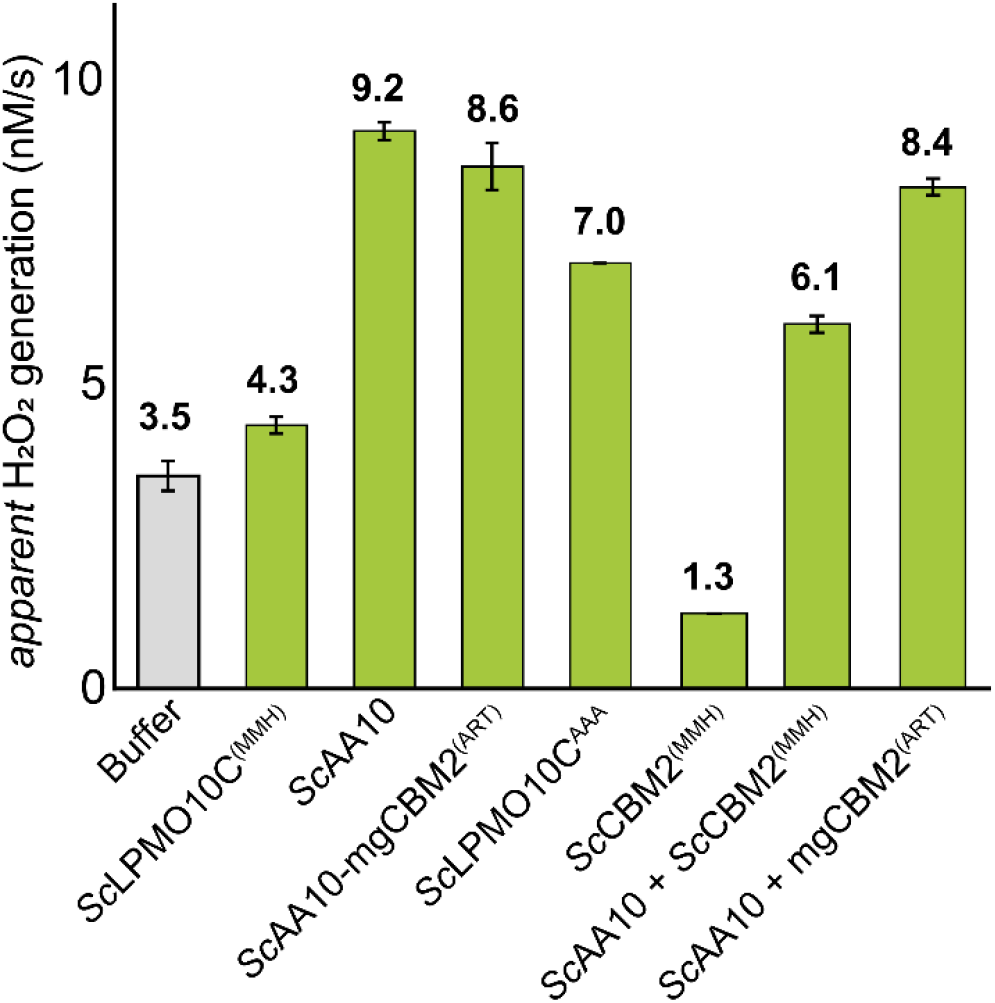
Oxidase activity of *Sc*LPMO10C variants measured using the Amplex Red/HRP assay. The bar chart shows the apparent rates of H_2_O_2_ production in various reactions containing various LPMOs or (combinations of) LPMO domains (green bars). A buffer control is shown for comparison (grey bar). All reactions were performed with 4 µM LPMO, with or without 4 µM CBM2, in 50 mM sodium phosphate buffer (pH 6.0). The reaction mixture contained 1 mM ascorbic acid, 5 U/mL HRP, 100 µM Amplex Red, and 1% (v/v) DMSO. Reaction rates were determined using the linear phase of the reaction (approximately 0–120 min), and error bars represent ± standard deviation (n = 3).

Interestingly, reactions with only *Sc*CBM2^(MMH)^ showed apparent oxidase activities that were lower than in control reactions with only buffer (Fig. 5). This again demonstrates that this CBM binds trace amounts of copper, that promote abiotic oxidation of the reductant. A similar effect was not observed for mgCBM2^(ART)^, which cannot bind copper (Fig. S8). As expected, based on the results discussed above, addition of *Sc*CBM2^(MMH)^ to reactions with *Sc*AA10 (Fig. 5) or mgAA10 (Fig. S8) reduced oxidase activity since free copper is sequestered from the reaction.

In a second assessment of copper reactivity, we determined the rate of reduction by ascorbic acid and reoxidation by H_2_O_2_ for full-length *Sc*LPMO10C^(MMH)^ and the catalytic domain only, *Sc*AA10, using fluorescence stopped-flow spectroscopy. Interestingly, while the two variants exhibited identical reduction rates (∼15,000 M^-1^ s^-1^), they showed differences in the rate of reoxidation of the Cu(I) state with H_2_O_2_ (10,700 ± 400 and 8400 ± 200 M^-1^ s^-1^, respectively; Fig. 6). This observation may be taken to suggest that, in the absence of cellulose substrate, the MMH-containing CBM interacts with the catalytic copper ion when it is in the Cu(I) state, thereby interfering with its potentially damaging reoxidation by H_2_O_2_.

**Figure 6.**
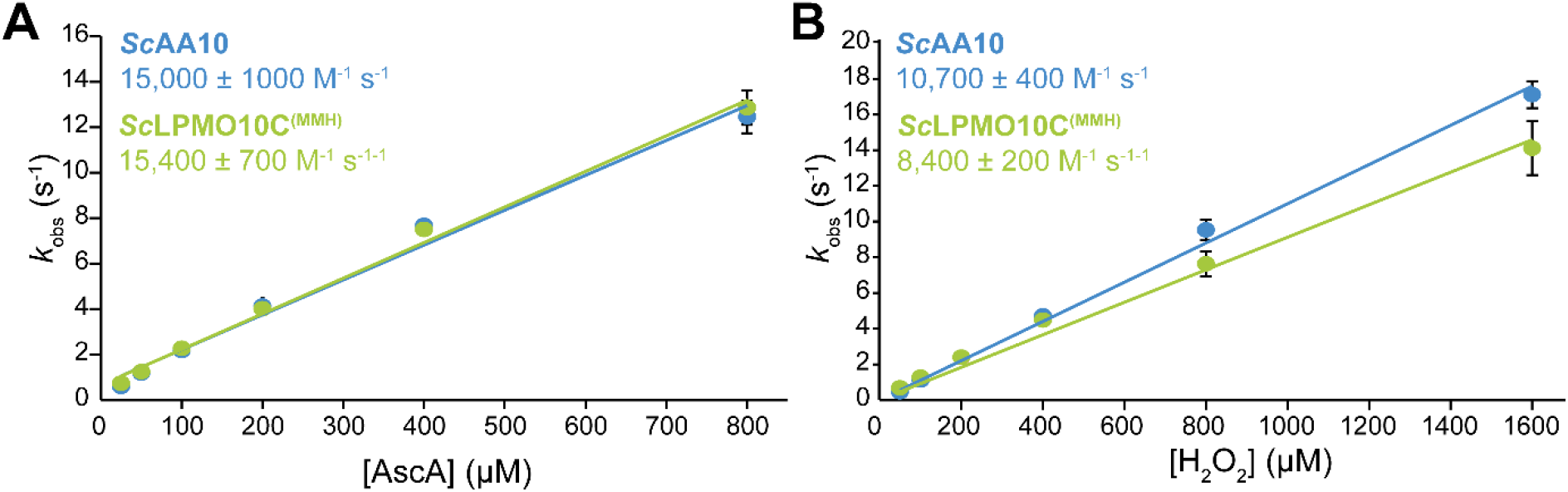
Kinetics of reduction and reoxidation of full-length *Sc*LPMO10C and its catalytic domain, *Sc*AA10. Panel A shows reduction by ascorbic acid (AscA) and panel B shows reoxidation by H_2_O_2_ under single-turnover conditions in the absence of substrate, both measured using fluorescence stopped-flow spectroscopy. The observed rate constants (*k*_obs_) were plotted against the concentration of AscA (A) or H_2_O_2_ (B) to determine the apparent second-order rate constants, which are shown in the graphs (*k*_AscA_ in A; *k*_H2O2_ in B). Error bars represent ±SD from three independent experiments (n = 3).

### Evaluating the effect of copper-binding CBMs on LPMO performance – Evidence for a CBM-CD interaction

Further proof for an interaction between the MMH-containing CBM2 and the reduced catalytic domain of the LPMO was obtained by experiments in which the LPMOs were placed under damaging conditions, namely reactions with reductant but lacking the substrate. As alluded to above, under such conditions, LPMO inactivation and release of copper from damaged active sites will occur, where the latter will lead to increased oxidation of ascorbic acid, higher levels of H_2_O_2_ and increasingly fast inactivation of the enzyme, in what is a self-reinforcing process [37]. Under these conditions, enzyme stability and the release of free copper may be observed by monitoring depletion of ascorbic acid [37, 36].

Fig. 7 shows that, expectedly, inactivation was fast, with AscA being fully depleted within the first 2–4 hours of the reaction, when a CBM with copper-binding properties was neither attached to, nor mixed with the catalytic domain. Two distinctly different reaction profiles emerged when a copper-binding CBM was included in the reactions. The first profile showed slow and steady consumption of AscA for up to 10–12 hours, as seen in reactions with wildtype *Sc*LPMO10C^(MMH)^ and mgAA10 with a CBM2 substitution (mgAA10-*Sc*CBM2^(MMH)^). This profile shows that, compared to the reactions without a copper-binding CBM2, copper is either released at a slower rate, due to a slower peroxidase reaction and reduced enzyme inactivation, or copper is released at the same pace but scavenged, where the former explanation entails an interaction between the CBM2 and the catalytic copper site, as also suggested by the data presented in Fig. 6B. The second reaction profile reflected an even slower rate of AscA consumption and was observed when combining a catalytic domain with a free MMH-containing CBM2 (Fig. 7A,B; note that AscA depletion for the rapidly inactivated catalytic domain is very fast). The difference between *Sc*LPMO10C^(MMH)^ and *Sc*AA10 *+ Sc*CBM2^(MMH)^, as well as the similar difference between mgAA10-*Sc*CBM2^(MMH)^) and mgAA10 + *Sc*CBM2^(MMH)^) has two possible explanations: either the free MMH-containing CBM2 has a higher capacity of scavenging free copper compared to the CBM bound to the CD, or inactivation and the resulting release of copper are slower for the enzymes with a covalently attached copper binding CBM. Both these explanations entail that the MMH-containing CBM interacts with the copper site in the CD, reducing the tendency of the CD to engage in damaging peroxidase reactions (relative to *Sc*AA10) and at the same time reducing the ability of the CBM2 to scavenge free copper (relative to *Sc*AA10 *+ Sc*CBM2^(MMH)^).

**Figure 7.**
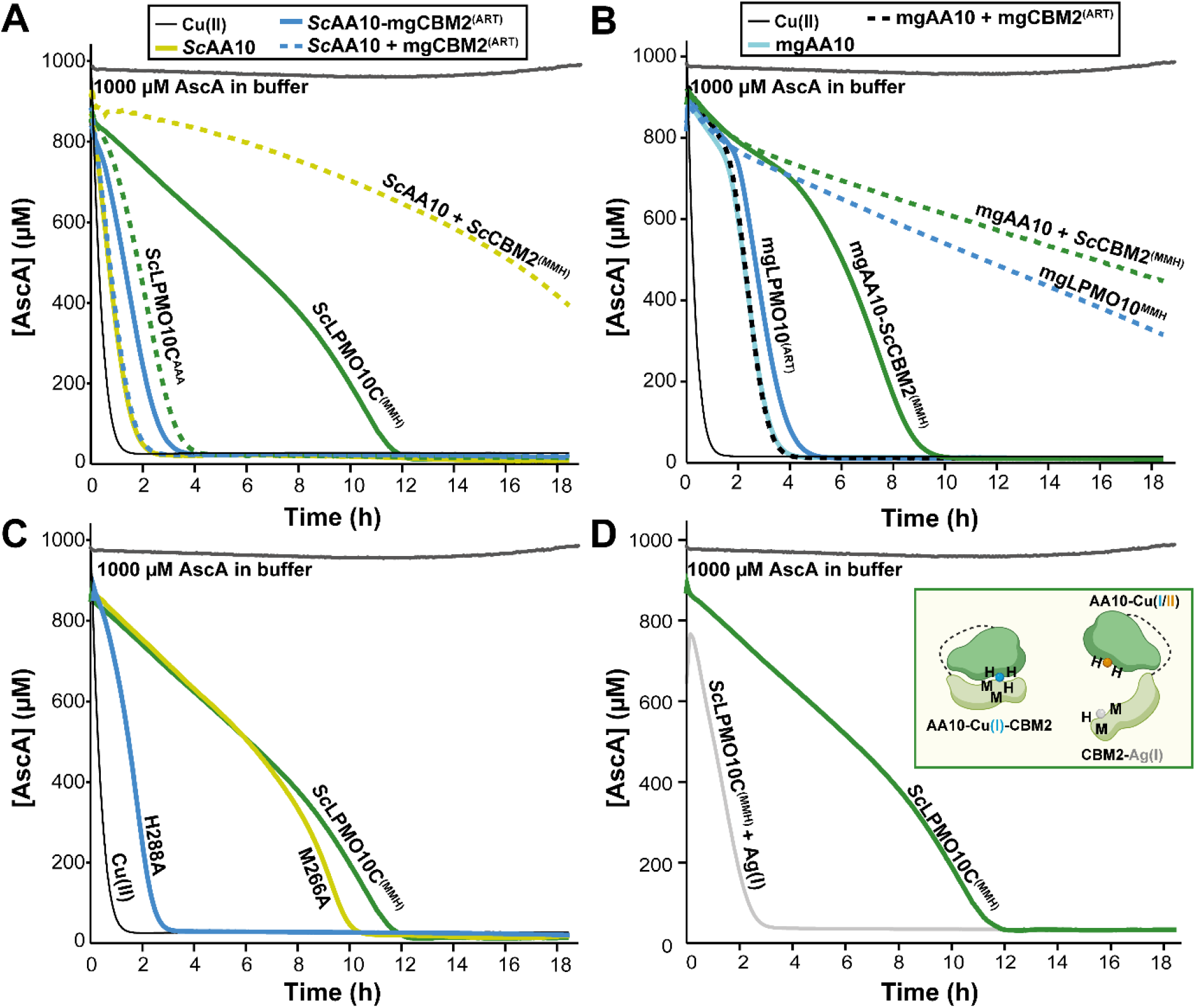
AscA depletion in wildtype and mutant LPMOs, alone and in combination with CBM2s displaying different copper-binding properties. The figure shows AscA depletion in cellulose-free reactions with *Sc*LPMO10C (A) and mgLPMO10 (B) variants. Panel C presents AscA depletion for single mutations in *Sc*LPMO10C (M266A and H288A), while panel D shows the effect of loading wildtype *Sc*LPMO10C^(MMH)^ with 3 µM Ag(I)NO_3_. The inset in panel D highlights that Ag(I)-binding CBM2s fail to interact with the catalytic domain. All reactions were performed with 1 µM LPMO, and in combination reactions, an additional 1 µM CBM2 (*Sc*CBM2^(MMH)^ or mgCBM2^(ART)^) was included. Reactions were carried out in 50 mM sodium phosphate (pH 6.0) with 1 mM AscA. A control reaction containing 1 µM Cu(II)SO_4_ was included to assess the effects of free copper, while a separate control with 1000 µM AscA in buffer (without LPMO) was used to evaluate AscA stability over time. Note that, in contrast to all other reactions reported in this study, these reactions were performed using (metal-free) TraceSelect water. Each reaction was conducted in technical triplicates (n=3); however, for clarity, only one representative curve per enzyme/combination is shown, as all replicates were essentially identical. Absorbance was measured at 255 nm every 3 min, and the AscA concentration was quantified using a standard curve of known AscA concentrations prepared in buffer. Reactions were incubated at 30 °C for up to 18 hours, with mixing occurring through plate movement during each measurement (i.e., every 3 minutes).

Interestingly, Fig. 7B shows that mgAA10-*Sc*CBM2^(MMH)^ shows the same inactivation behavior as *Sc*LPMO10C^(MMH)^, whereas mgLPMO10^MMH^ shows the same very slow inactivation kinetics as in reactions with a separate CD and copper-binding CBM2. This suggests that the mgCBM2^MMH^ domain does not interact as well with mgAA10 as does the *Sc*CBM2^(MMH)^ domain, allowing it to scavenge free copper as if it were moving free in solution. This observation supports the notion derived from the AlphaFold predictions (see above) that other structural features beyond the MMH motif affect the interaction between the CD and the CBM2.

Additional experiments with single mutants in full-length *Sc*LPMO10C, showed that mutation of the His in the MMH motif (H288A), which is the residue involved in the predicted interdomain interaction, abolished the protective function of the CBM2 (rapid inactivation, similar to that of the CD only). On the other hand, mutation of one of the copper-binding methionines (M266A), which do not seem to be involved in the interdomain interaction, had only marginal effects on the redox stability of the full-length enzyme (Fig. 7C). These effects were also reflected in cellulose oxidation experiments (Fig. S9A).

Using Cu(I) binding proteins with methionine motifs such as CusF [57], it has been shown that, due to structural similarities in ligand preferences, coordination geometry, and ionic radius, Cu(I) can be replaced by silver [Ag(I)], which is not redox-active. As a final verification of Cu(I)-binding by *Sc*CBM2^(MMH)^, we pre-saturated wildtype *Sc*LPMO10C with AgNO_3_ prior to performing the AscA depletion assay (Fig. 7D) and measuring cellulose oxidation (Fig. S9B). An Ag(I)-binding CBM would be unable to interact with the active site copper or chelate released copper, which should lead to earlier inactivation, and, thus, more rapid depletion of AscA. Fig. 7D shows that, indeed, inactivation happened considerably earlier in reactions with the Ag(I) pre-loaded enzyme compared to reactions without added Ag(I). The Ag(I) loaded enzyme shows a depletion curve close to the one observed for the CD alone (Fig. 7A). In cellulose oxidation experiments the Ag(I)-loaded enzyme showed a higher initial rate (Fig. S9B), exactly as was observed for *Sc*LPMO10C^AAA^ and *Sc*AA10-mgCBM2^(ART)^ (Fig. 4), which lack the copper site.

### Related copper sites in CBM2s

Through our investigation, we identified additional potential copper sites (similar to the MMH site) in the CBM2s of other LPMOs, though with some variation in the ligands and relative positions of these sites (Fig. S10). One such example is *Af*LPMO10B, a previously uncharacterized enzyme, which we produced and characterized. Phylogenetic analysis and assessment of enzyme activity (Fig. S11) place *Af*LPMO10B in subclade A1 within clade II, together with *Sc*LPMO10C, mgLPMO10 and other cellulose-active C1-oxidizing enzymes. The putative copper-binding site in the CBM2 of this LPMO contains two histidines (His298 and His350) and two methionines (Met280 and Met296), and AlphaFold predicts that both histidines interact with the catalytic copper (Fig. 8A). In line with the predictions made for variants of *Sc*LPMO10C and mgLPMO10 and their CBM2s, AlphaFold did not predict such an interaction in the absence of copper (Fig. 8B) and predicted that the isolated CBM2 binds copper (Fig. 8C). We produced the full-length enzyme and the CBM2, *Af*CBM2^(MMHH)^.

**Figure 8.**
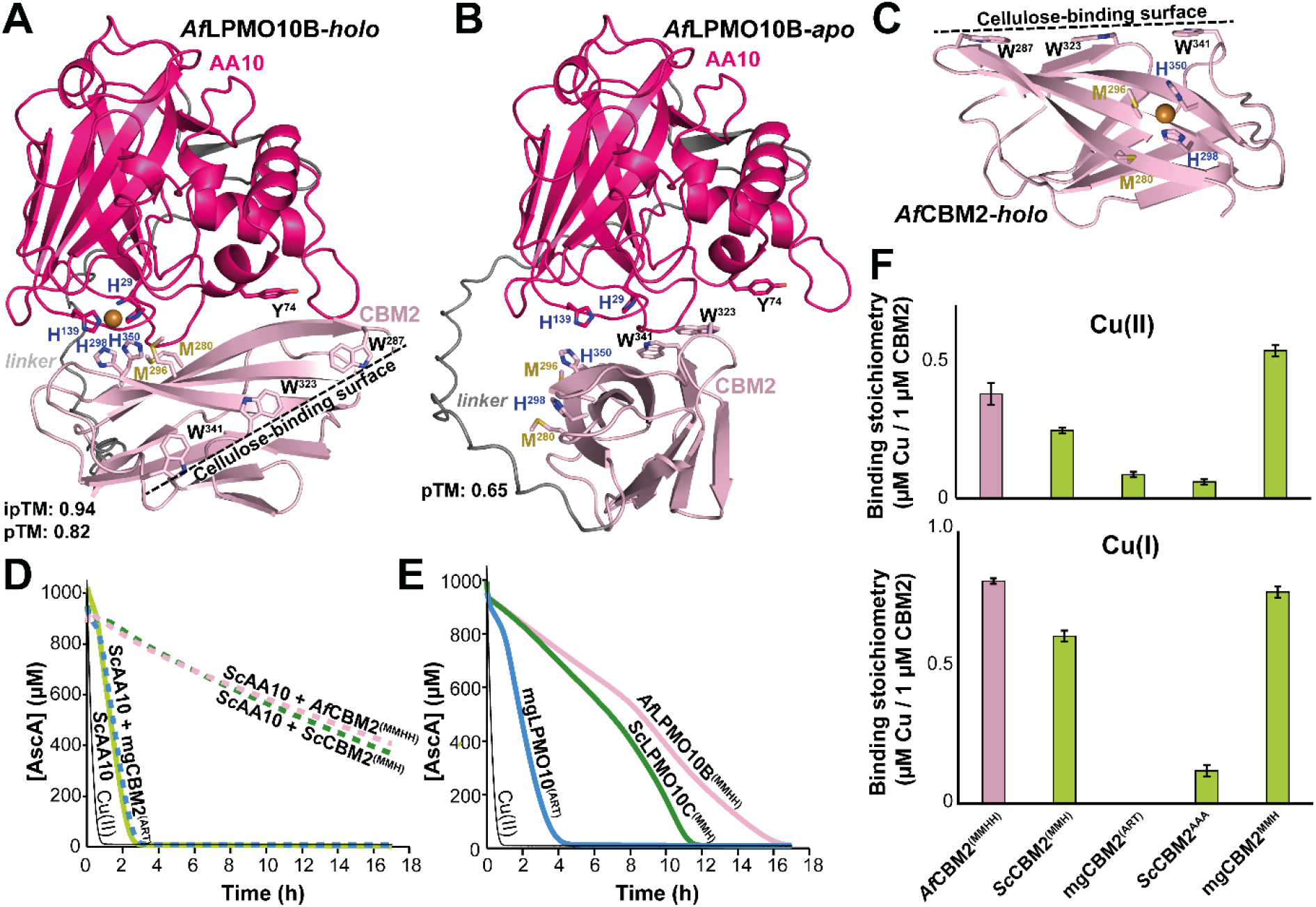
Structural prediction and experimental analysis of copper binding in *Af*LPMO10B^(MMHH)^ and its implications for enzyme stability. The predicted AlphaFold3 structures of *holo* (A) and *apo* (B) *Af*LPMO10B^(MMHH)^, along with copper interactions in the isolated CBM2 (C), are similar to those obtained for *Sc*LPMO10C^(MMH)^ and its CBM2 (Fig. 1). Panel D displays enzyme inactivation monitored using the AscA depletion assay for *Sc*AA10 in the presence of three wildtype CBMs with ART, MMH, and MMHH motifs. Panel E shows inactivation curves for the three full-length enzymes investigated in this study: *Af*LPMO10B^(MMHH)^, mgLPMO10^(ART)^, and *Sc*LPMO10C^(MMH)^. Reactions were conducted in 50 mM sodium phosphate (pH 6.0) with 1 mM AscA, and each reaction was performed in triplicates (n=3); for clarity, only one representative curve per enzyme/combination is shown, as all replicates were essentially identical. Panel F summarizes the results of the copper binding assays for all five CBM2s (three wildtypes and two mutants) for Cu(II) (*upper*) and Cu(I) (*lower*) binding. Reactions were carried out as described in the legend of Fig. 3 using 4 µM CBM and 8 µM Cu(II)SO_4_, with and without AscA.

The inactivation behavior of *Af*LPMO10B in the ascorbic acid depletion assay was similar to that of *Sc*LPMO10C, displaying slow consumption of ascorbate for the full-length enzyme (Fig. 8E). When the *Sc*AA10 catalytic domain was mixed with *Af*CBM2^(MMHH)^, AscA depletion became even slower, indicative of copper chelation and similar to what was observed in reactions with *Sc*CBM2^(MMH)^, but not mgCBM2^(ART)^ (Fig. 8D). Finally, copper binding assays using the BCS-Cu(I) fluorescence assay demonstrated that *Af*CBM2^(MMHH)^ exhibits a clear affinity for Cu(I) and, to a lesser extent, Cu(II) (Fig. 8F).

## Concluding Remarks

The discovery of CBM2s with copper affinity that have likely co-evolved with LPMOs and that influence the activity and stability of these enzymes, adds a new dimension to our understanding of the roles of CBMs in Nature. Our results suggest a dual role for these CBM2s, which bind copper through a conserved MMH motif: they can bind copper in solution, preferably in the Cu(I) state, and they modulate the reactivity of the reduced catalytic copper by interacting with it. The latter interaction is predicted by AlphaFold and is supported by several of the experimental observations described above. One indication is the reduced reoxidation rate observed for *Sc*LPMO10C^(MMH)^ compared to *Sc*AA10. Another indication is the slower depletion of AscA in substrate-free reactions with *Sc*AA10 *+ Sc*CBM2^(MMH)^ compared to reactions with wildtype *Sc*LPMO10C^(MMH)^. Finally, we show that the presence of the MMH motif in the CBM2 of *Sc*LPMO10C reduces the initial rate of the reaction with cellulose, suggesting an interaction between this motif and the catalytic copper. It is worth noting that the copper site in the LPMO-CBM2 complex, with its additional His ligand relative to the LPMO alone (Fig. 1) shows a structural arrangement similar to that of the Cu(B) site in pMMOs (Fig. S12).

From a biological perspective, the present findings shed light on how LPMOs may operate in natural environments, particularly with regard to maintaining redox stability under oxidative stress conditions, such as substrate depletion or elevated H_2_O_2_ levels. The MMH containing CBM2 prevents the self-reinforcing cycle of oxidative damage and enzyme inactivation. Importantly, the fact that the copper binding site and the cellulose-binding residues are located on different faces of the CBM2 (Fig. 1) suggests an appealing, albeit speculative regulatory scenario. In the absence of substrate, the CBM2 is likely to bind to the reduced catalytic copper, preventing it from engaging in off-pathway reactions. On the other hand, one could envisage that binding of the CBM2 to its substrate leads to a conformational change that abolishes the interaction of its MMH site with the catalytic copper (Fig. 9). In that way, the copper site would become available for catalytic action only when the substrate is near, which is consistent with a biologically relevant regulatory mechanism.

**Figure 9.**
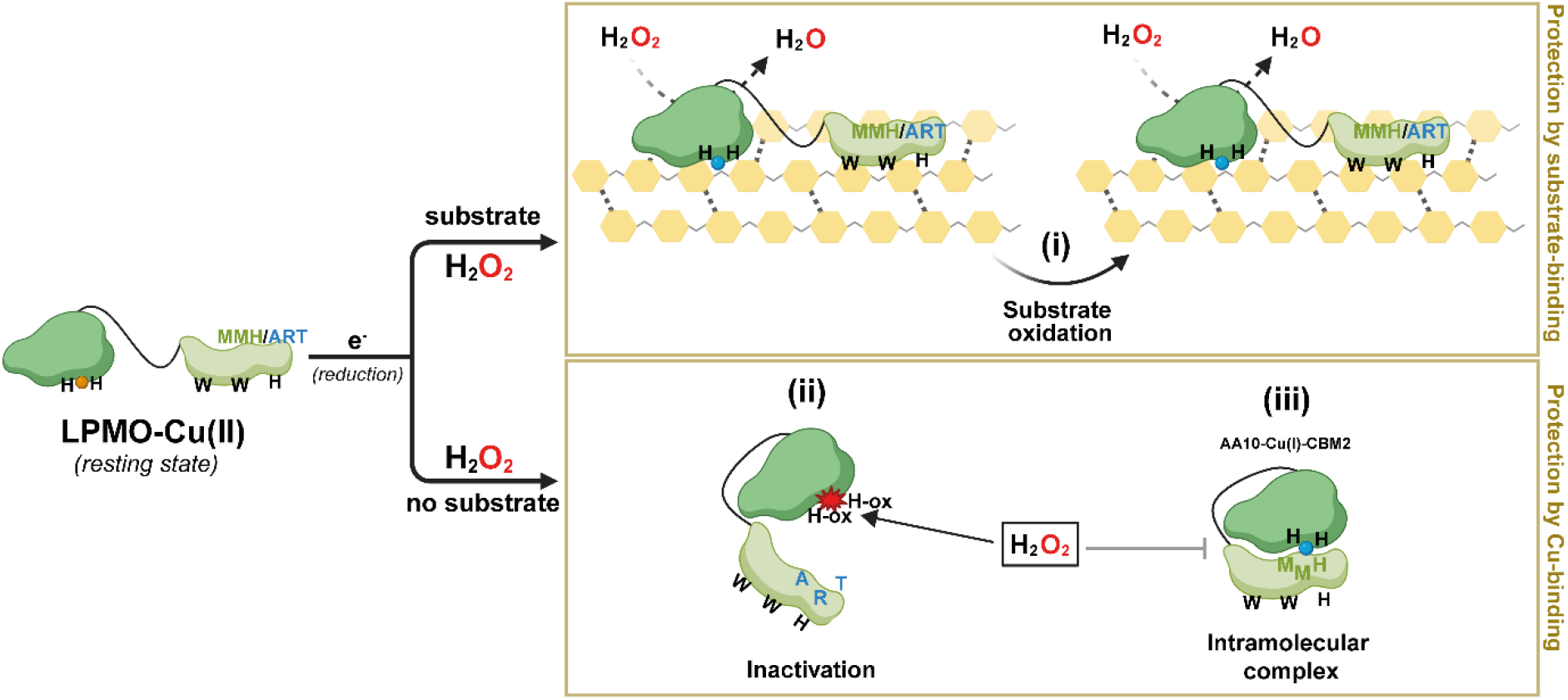
Proposed mechanism for regulation of copper reactivity in LPMOs by appended CBM2s. Schematic overview of the possible reactions following reduction of an AA10-CBM2 enzyme in the presence or absence of substrate. In the presence of substrate, the enzyme exhibits productive peroxygenase activity. The carbohydrate-binding module (CBM2) enhances substrate affinity, ensuring H_2_O_2_ resulting from oxidation of the reductant is used productively, thus protecting the enzyme from damaging off-pathway reactions, regardless of the presence of the MMH copper site (i). In the absence of substrate, the reduced LPMO is prone to engaging in damaging peroxidase reactions, leading to rapid enzyme inactivation (ii). When the CBM has evolved to interact with the catalytic domain through its MMH copper site, perhaps facilitated by other structural features of the MMH-containing face of the CBM, the enzyme adopts a closed conformation, akin to a clam, reducing copper reactivity and, thus enzyme inactivation (iii). Importantly, the copper site and cellulose-binding surface lie on opposite faces of the CBM2, suggesting that substrate binding triggers the opening of the “clam,” relieving CBM-mediated inhibition of LPMO activity and enabling productive catalysis in the presence of substrate.

As alluded to above, copper is an important transition metal, the levels of which need to be carefully regulated in living species and ecosystems [2, 7, 8]. Methionines are well-known as critical residues in copper trafficking proteins (e.g., CusF and CopC), where they provide binding sites that facilitate metal transfer. The copper-binding CBMs identified here may serve a similar role by scavenging copper from the environment and delivering it to the catalytic center of newly synthesized *apo*-LPMOs.

Next to revealing a completely new function of CBMs, not directly related to carbohydrate-binding, the present findings shed light on the functional implications of modularity in copper-containing LPMOs and, potentially, other metalloenzymes. Quite some LPMOs are associated with a wide variety of modules whose functions remain largely unexplored. It is tempting to speculate that such appended domains may have evolved to stabilize these powerful redox enzymes under conditions that promote destabilizing off-pathway reactions, as we show here for CBM2 containing *Sc*LPMO10C and *Af*LPMO10B. The newly discovered role of copper-binding CBM2 domains, which perhaps can be extended to other domains appended to LPMOs, provides novel biological insight into how enzyme activity and stability are regulated, and may eventually help in the engineering of better LPMOs for optimized industrial application.

## Materials and Methods

### Comprehensive screening of CBM2 domains for MMH motifs

The complete set of proteins containing CBM2 domains (matching the InterPro entry IPR001919; 26,436 sequences) was retrieved from the InterPro database [72]. Duplicate sequences were removed using CD-HIT [73], resulting in 25,466 unique CBM2-containing proteins. The subsequent steps of the analysis were carried out using an in-house developed Python script, available in the notebook CBM2_MMH_analysis.ipynb (github.com/EstebanLT/CBM2_analysis). The CBM2 domains were extracted from the full protein sequences and the motif MX_n_MXH (MMH) was searched using the regular expression pattern “M.*M.H”, resulting in 713 CBM2 sequences containing the MMH motif. The structures of these proteins were retrieved from the AlphaFold database [64, 74], and the ones missing were predicted using the public Colabfold batch notebook [75] or using the AlphaFold3 server [51]. The presence of MMH motifs was confirmed in 649 LPMO sequences by automatically identifying two methionines and a histidine within 8.5 Å (Cα-Cα distance) using the Python script described above, followed by manual inspection of special cases in PyMOL. An additional 34 protein sequences were found among the AlphaFold-verified MMH-containing sequences but were not annotated as LPMOs according to the dbCAN3 server [76]. To assess whether the MMH-containing AA10 sequences also featured a canonical LPMO REF motif, the same Python script was used to search for the RX_3-10_EXF (REF) motif within their AA10 domains using regular expressions.

### Sequence space and phylogenetics analyses

The dbCAN3 server was used to fetch all annotated AA10 LPMO sequences (database 07262023). CD-HIT [73] was used to remove duplicate sequences, reducing the dataset to unique AA10 LPMOs. MAFFT [77] (FFT-NS-1 option) and fasttree [78] (default options) were used to obtain a phylogenetic tree. Taxonomic coloring of the tree was achieved using phylo-color.py (https://github.com/acorg/phylo-color). A subset of all CBM2-containing LPMOs was used to analyze the occurrence of the MMH motif and its relation to the 2^nd^ coordination sphere of the copper in the catalytic domains. This subset of sequences was aligned with MAFFT (L-INS-I) to perform the motif search in the sequence space. For this purpose, the in-house script count-aminoacid-combinations.py (github.com/IAyuso) was used, using as target M266, M286 and H288 from the CBM2 present in *Sc*LPMO10C. To investigate correlations with the catalytic domain, we focused on Arg212, Glu217, and Phe219 (*Sc*LPMO10C numbering) as key residues in the second coordination sphere of the copper center in AA10 LPMOs. Our in-house script reports all possible combinations of these positions in the alignment, as found in the dataset, along with the corresponding sequence names. Using this approach, we identified AA10 LPMOs containing REF, MMH, or REF-MMH motifs, which were subsequently mapped and visualized in the phylogenetic tree.

### Cloning, mutagenesis, and CBM substitutions

Two well-characterized bacterial LPMOs were selected for cloning and mutagenesis to introduce or remove predicted copper-binding residues (see Table 1). These include LPMO10C from *Streptomyces coelicolor* A3(2), referred to as *Sc*LPMO10C (UniProtKB: Q9RJY2), and a metagenome-derived LPMO designated mgLPMO10, which was identified as highly expressed in a rice straw-derived metatranscriptome [79]. The full-length wildtype sequences of these enzymes, as well as their isolated catalytic domains, were previously cloned into the pRSETB expression vector [52, 65], which includes the signal peptide from *Sm*LPMO10A to ensure efficient secretion of the mature (signal peptide-free) protein into the periplasm [10]. A third LPMO, *Af*LPMO10B from *Actinoplanes friuliensis* DSM 7358 (UniProtKB: U5W274) was cloned in the same manner. The CBM2s from all three LPMOs were also cloned individually, each fused downstream of the *Sm*LPMO10A signal peptide to enable periplasmic expression. The regions cloned were as follows: residues 275–373 for *Af*CBM2, 265–363 for mgCBM2, and 263–364 for *Sc*CBM2.

Plasmids encoding full-length *Sc*LPMO10C and mgLPMO10, as well as their corresponding CBM2s, were used as templates for site-directed mutagenesis. Mutants listed in Table 1 were generated by sequentially introducing one or two mutations using the QuikChange II XL site-directed mutagenesis kit (Agilent Technologies).

CBM2 domain substitutions in *Sc*LPMO10C and mgLPMO10 were carried out via inverse PCR. The plasmids were amplified excluding the CBM2-encoding region, while the respective CBM2 domains were amplified in parallel using primers with overhangs matching the plasmid ends. The amplified CBMs were inserted into the plasmids using the In-Fusion® HD Cloning Kit (Clontech).

All LPMO sequences used in this study were codon-optimized for *Escherichia coli* expression. All newly generated plasmids were verified by sequencing prior to transformation into One Shot® BL21 Star™ (DE3) chemically competent *E. coli* cells, for subsequent expression and protein production and purification.

### LPMO production and purification

Cells harboring the expression plasmids were inoculated into lysogeny broth (LB) supplemented with 100 µg/mL ampicillin. All variants (wildtypes and mutants), with the exception of *Sc*AA10, were cultivated at 37 °C for approximately 20 hours. Cells producing *Sc*AA10 were instead grown at 30 °C for 24 hours to enhance soluble expression. Protein expression was driven by basal (leaky) activity of the T7 promoter, and no isopropyl β-D-1-thiogalactopyranoside (IPTG) was added.

Cells were harvested by centrifugation, and periplasmic extracts were prepared using an osmotic shock method as previously described [80]. The resulting periplasmic fractions, containing mature (i.e., signal peptide-free) proteins, were sterilized by filtration prior to purification. All *Af*LPMO10B and *Sc*LPMO10C variants, along with full-length mgLPMO10 variants, including those with CBM substitutions, were purified by anion-exchange chromatography using a 5 mL HiTrap DEAE FF column (Cytiva, MA, USA) equilibrated with 50 mM Tris-HCl (pH 7.5). Bound proteins were eluted with a linear NaCl gradient (0–500 mM) applied over 60 column volumes.

Truncated mgLPMO10 variants (mgAA10 and mgCBM2^ART/MMH^) were first purified by hydrophobic interaction chromatography using a 5 mL HiTrap Phenyl FF column (Cytiva, MA, USA) with a running buffer of 50 mM Tris-HCl (pH 8.0) containing 1 M ammonium sulfate. Elution was achieved by decreasing the salt concentration from 1 M to 0 M over 25 column volumes.

LPMO-containing fractions were identified by SDS-PAGE, pooled, and concentrated using Amicon® Ultra centrifugal filters (Millipore, Darmstadt, Germany) with molecular weight cutoffs of 10 kDa for catalytic domains and full-length proteins, or 3 kDa for CBMs. Final purification was performed using preparative size-exclusion chromatography on a ProteoSEC Dynamic 16/60 3–70 HR column (Protein Ark, Sheffield, UK), operated at 1 mL/min in 50 mM Tris (pH 7.5) containing 200 mM NaCl. Protein purity was confirmed by SDS-PAGE, and fractions containing pure protein were pooled and concentrated as described above.

### Copper and silver saturation

Purified LPMO variants (excluding the isolated CBM2s) were incubated with a two-fold molar excess of CuSO_4_ at room temperature for 30 minutes to ensure copper saturation. Unbound copper was removed by multiple rounds of buffer exchange via dilution and concentration using Amicon® Ultra centrifugal filters and 50 mM sodium phosphate buffer (pH 6.0). The cumulative dilution factor achieved during this process exceeded 1,000,000, effectively eliminating free copper ions from the samples.

To bind Ag(I) to the CBM2, Ag(I)NO_3_ was added to the copper-saturated enzyme in buffer at either a 1- or 3-fold molar ratio, 30 minutes prior to initiating the LPMO reaction by adding substrate and ascorbic acid.

### Cellulose binding

The equilibrium binding constants (*K*_d_) and binding capacities (*B*_max_) for wildtype mgCBM2 and *Sc*CBM2 were determined by incubating protein solutions at varying concentrations (0, 10, 25, 50, 75, 150, 300, and 500 μg/mL) with 10 g/L Avicel. Prior to Avicel addition, the A_280_ was measured for each protein solution (in 50 mM sodium phosphate buffer, pH 6.0) to generate individual standard curves for each protein variant. After adding Avicel, the solutions were incubated at 22 °C with agitation (800 rpm) in an Eppendorf Thermomixer C for 60 min. The samples were then filtered using a 96-well filter plate (Millipore, Darmstadt, Germany), and the concentration of unbound protein in the supernatant was measured by A_280_. All assays were performed in triplicate with blanks (buffer and 10 g/L Avicel). The equilibrium dissociation constant (*K*_d_, μM) and substrate binding capacity (*B*_max_, μmol/g Avicel) were calculated by fitting the binding data to the one-site binding equation [*P*_bound_] = *B*_max_ [*P*_free_] / (*K*_d_ + [*P*_free_]), using nonlinear regression in GraphPad Prism 10 (GraphPad, CA, USA).

### Copper binding (BCS) assay

To assess copper binding by LPMOs, 4 µM protein was incubated with 8 µM Cu(II)SO_4_ in 50 mM sodium phosphate buffer (pH 6.0) for 10 minutes at room temperature in the presence or absence of 20 µM ascorbic acid. Next, CBMs were removed from the solutions using microcentrifugal filters with a 3 kDa molecular weight cut-off (VWR, Radnor, PA, USA; catalog number 82031-346; 5 minutes at 10,000 × g) and 100 µL filtrates were transferred into non-transparent (black) 96-well microtiter plates (Thermo Fisher Scientific, MA, USA). The residual (i.e., unbound) copper in filtrates was quantified using bathocuproine disulfonate (BCS), a Cu(I)-specific probe which displays a decrease in fluorescence upon copper binding [67, 68]. 100 µL of 40 µM BCS solution in 50 mM sodium phosphate buffer (pH 6.0) containing 40 µM ascorbic acid was mixed with each free copper sample and incubated for 10 minutes at room temperature inside a Varioskan LUX plate reader (Thermo Fisher Scientific, MA, USA). Fluorescence was measured every minute (λ_Ex/Em_ = 290/325 nm) during the incubation to ensure signal stability. The final (10-minute) measurements were used to calculate free copper concentration according to a standard curve made with Cu(II)SO_4_ solutions (0 – 8 µM) treated exactly the same way as experimental samples.

### Inductively coupled plasma mass spectrometry (ICP-MS)

ICP-MS was employed to quantify the copper content in selected *Sc*LPMO10C and mgLPMO10 variants. Copper-saturated LPMOs (2 µM) in TraceSELECT™ (Honeywell, Charlotte, NC, USA) water were analyzed using an Agilent 8800 ICP-QQQ tandem quadrupole mass spectrometer (Agilent Technologies, Santa Clara, CA, USA). The instrument was operated in triple quadrupole mode, utilizing helium as the collision gas in the collision/reaction cell to reduce diatomic interferences originating from the plasma or sample. Prior to analysis, samples were mixed with a multi-element internal standard (Inorganic Ventures, Christiansburg, VA, USA) and 65% NORMAPUR® nitric acid (VWR, Radnor, PA, USA), then autoclaved at 121°C for 30 minutes in sealed tubes. Following cooling, the samples were diluted with 18.2 MΩ type I deionized water to achieve a final 10% (v/v) nitric acid concentration. Instrumental drift was monitored and corrected by analyzing a control standard (Inorganic Ventures) between samples. Calibration curves were generated prior to sample measurements, and copper concentrations were determined using indium as an internal standard. All experiments were conducted in duplicate (n = 2) to ensure reproducibility.

### Cellulose degradation

LPMO reactions with insoluble substrate (1% w/v Avicel) were performed in 50 mM sodium phosphate buffer (pH 6.0) at 40 °C and 800 rpm using an Eppendorf Thermomixer® C (Eppendorf, Hamburg, Germany). Unless otherwise specified, reactions contained 1 µM LPMO, either alone or in combination with 1 µM CBM2, and were initiated by the addition of 1 mM ascorbic acid (final concentration). At various time points, aliquots were collected and the reactions were quenched by removing the insoluble substrate using a 96-well filter plate (Millipore, MA, USA). The resulting filtrates were treated with 1 µM recombinant *Thermobifida fusca* GH6 endoglucanase (*Tf*Cel6A; produced in-house) [81] and incubated statically at 37 °C overnight to convert soluble C1-oxidized oligosaccharides into a simple mixture of oxidized dimers and trimers (GlcGlc1A and Glc_2_Glc1A).

### Product quantification

High-performance anion-exchange chromatography with pulsed amperometric detection (HPAEC-PAD) of cellulose-derived products was performed as described previously [82], using an ICS-5000 system (Thermo Fisher Scientific, Waltham, MA, USA) equipped with a disposable electrochemical gold electrode. Samples (5 µL) were injected onto a CarboPac PA200 column (3 × 250 mm), operated with 0.1 M NaOH (eluent A) at a flow rate of 0.5 mL/min and a column temperature of 30 °C. Elution was achieved using a stepwise gradient with increasing concentrations of eluent B (0.1 M NaOH + 1 M NaOAc) as follows: 0–5.5% B over 3 min; 5.5–15% B over 6 min; 15–100% B over 11 min; 100–0% B over 0.1 min; and 0% B (reconditioning) for 5.9 min.

For samples not treated with *Tf*Cel6A, a more gradual elution profile was used: 0–5.5% B over 4.5 min; 5.5–15% B over 9 min; 15–100% B over 16.5 min; 100–0% B over 0.1 min; and 0% B (reconditioning) for 8.9 min. Chromatograms were recorded and analyzed using Chromeleon 7.0 software (Thermo Fisher Scientific, Waltham, MA, USA). LPMO products were quantified using standard mixtures of C1-oxidized cellobiose and cellotriose (GlcGlc1A and Glc_2_Glc1A), which were produced in-house using cellobiose dehydrogenase, according to a previously published protocol [83].

### Oxidase activity assay and hydrogen peroxide detection

Generation of hydrogen peroxide (H_2_O_2_) was measured using a modified HRP/Amplex Red assay [66], based on the protocol by Kittl et al. [69]. Briefly, 90 µL of LPMO solution (4 µM LPMO in the final reaction), with or without the addition of 4 µM CBM2, in 50 mM sodium phosphate buffer (pH 6.0), containing horse radish peroxidase (HRP) and Amplex Red, was pre-incubated for 5 min at 30 °C in a 96-well microtiter plate. The reaction was initiated by adding 10 µL of 10 mM ascorbic acid, followed by 10 s of mixing in a Varioskan LUX plate reader (Thermo Fisher Scientific, MA, USA). The final concentrations in the reactions were 4 µM LPMO ± 4 µM CBM2, 5 U/mL HRP, 100 µM Amplex Red and 1 mM AscA. H_2_O_2_ production was monitored by measuring resorufin formation at 540 nm. Standards with known concentrations of H_2_O_2_ were prepared in 50 mM sodium phosphate buffer (pH 6.0) with 1 mM ascorbic acid, followed by the addition of HRP and Amplex Red. Non-enzymatic H_2_O_2_ production was evaluated in control reactions containing 1 mM ascorbic acid and buffer (i.e., without LPMO).

### Stopped-Flow redox kinetics

Fluorescence differences between the LPMO-Cu^2+^ and LPMO-Cu^+^ forms were used to monitor the kinetics of reduction by ascorbate and oxidation by H_2_O_2_, as previously described [83, 31]. All experiments were performed using a SFM-4000 stopped-flow equipped with a MOS 200M dual absorbance fluorescence spectrometer (BioLogic, Seyssinet-Pariset, France). The photomultiplier tube was equipped with a 340 nm bandpass filter and the high voltage was set to 600 V. The excitation wavelength was 280 nm, and fluorescence was collected using a 340 nm bandpass filter, with an increase observed during reduction and a decay during oxidation. Experiments were conducted at 25 °C in 50 mM sodium phosphate (pH 6.0). To monitor reduction of Cu^2+^ to Cu^+^, LPMO-Cu^2+^ (5 µM of *Sc*AA10 or *Sc*LPMO10C^MMH^) was mixed with varying concentrations of ascorbate (25–800 µM final concentration). For oxidation, a double-mixing stopped-flow approach was used. First, the LPMO-Cu^2+^ (10 µM initial concentration) was mixed with one molar equivalent of ascorbate for 10 seconds to form LPMO-Cu^+^. Secondly, the *in situ* generated LPMO-Cu^+^ was mixed with H_2_O_2_ (50–1600 µM after mixing), and fluorescence decay was monitored. Prior to setting up the instrument, all stock solutions were deoxygenated by N_2_ sparging before storage in an A95TG anaerobic workstation (Don Whitley Scientific, West-Yorkshire, UK) for at least 16 hours. Working dilutions were prepared in sealed syringes inside an anaerobic chamber. Before all experiments, the stopped-flow system was thoroughly flushed with deoxygenated buffer to maintain anaerobic conditions. The observed rate constants (*k*_*obs*_) were determined by solving a single exponential equation 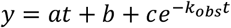. The *k*_*obs*_ values were plotted against ascorbate or H_2_O_2_ concentrations and fitted using linear least-squares regression to determine the apparent second-order rate constants (k_app,_^AscA^ for reduction and k_app,_^H2O2^ for oxidation), assuming pseudo-first-order conditions. All experiments were conducted in triplicates.

### Ascorbic acid depletion

Reactions containing 2 µM LPMO, either alone or in combination with 2 µM CBM2, were preincubated in 50 mM sodium phosphate buffer (pH 6.0) at 30 °C for 5 min. After preincubation, 50 µL of the protein solution was mixed with 50 µL of a 2 mM ascorbic acid (AscA) solution directly in a 96-well UV-transparent plate (Corning, Corning, NY, USA), resulting in final protein concentrations of 1 µM and 1 mM AscA. AscA depletion was monitored spectrophotometrically by measuring absorbance at 255 nm every 3 min for up to 19 h using a Varioskan LUX plate reader (Thermo Fisher Scientific, Waltham, MA, USA). AscA concentrations were determined using a standard curve generated from known concentrations of AscA measured at 255 nm. All experiments were performed in triplicates (n = 3). All solutions used in these experiments were prepared with TraceSELECT™ water.

## Acknowledgements and Funding Sources

We would like to thank MSc student Kamilla Løvheim Kleppang for cloning and purification of *Af*LPMO10B and *Af*CBM2. We gratefully acknowledge funding from the European Research Council (ERC) through a Synergy Grant (856446). We also acknowledge the Research Council of Norway for funding equipment through grants 245828 and 270038.

## Notes

### Competing Interest Statement

The authors have declared no competing interest.

## References

1. B. Halliwell, J. M. C. Gutteridge, Oxygen toxicity, oxygen radicals, transition metals and disease. Biochem. J. 219, 1–14 (1984).

2. A. K. Boal, A. C. Rosenzweig, Structural biology of copper trafficking. Chem. Rev. 109, 4760–4779 (2009).

3. T. D. Rae, P. J. Schmidt, R. A. Pufahl, V. C. Culotta, T. V. O’Halloran, Undetectable intracellular free copper: the requirement of a copper chaperone for superoxide dismutase. Science 284, 805–808 (1999).

4. A. Changela, K. Chen, Y. Xue, J. Holschen, C. E. Outten et al., Molecular basis of metal-ion selectivity and zeptomolar sensitivity by CueR. Science 301, 1383–1387 (2003).

5. L. A. Finney, T. V. O’Halloran, Transition metal speciation in the cell: insights from the chemistry of metal ion receptors. Science 300, 931–936 (2003).

6. B. E. Kim, T. Nevitt, D. J. Thiele, Mechanisms for copper acquisition, distribution and regulation. Nat. Chem. Biol. 4, 176–185 (2008).

7. R. A. Festa, D. J. Thiele, Copper: an essential metal in biology. Curr. Biol. 21, R877–R883 (2011).

8. J. T. Rubino, K. J. Franz, Coordination chemistry of copper proteins: how nature handles a toxic cargo for essential function. J. Inorg. Biochem. 107, 129–143 (2012).

9. R. J. Quinlan, M. D. Sweeney, L. Lo Leggio, H. Otten, J. C. Poulsen et al., Insights into the oxidative degradation of cellulose by a copper metalloenzyme that exploits biomass components. Proc. Natl. Acad. Sci. U. S. A. 108, 15079–15084 (2011).

10. G. Vaaje-Kolstad, B. Westereng, S. J. Horn, Z. Liu, H. Zhai et al., An oxidative enzyme boosting the enzymatic conversion of recalcitrant polysaccharides. Science 330, 219–222 (2010).

11. Z. Forsberg, G. Vaaje-Kolstad, B. Westereng, A. C. Bunæs, Y. Stenstrøm et al., Cleavage of cellulose by a CBM33 protein. Protein Sci. 20, 1479–1483 (2011).

12. C. M. Phillips, W. T. Beeson, J. H. Cate, M. A. Marletta, Cellobiose dehydrogenase and a copper-dependent polysaccharide monooxygenase potentiate cellulose degradation by Neurospora crassa. ACS Chem. Biol. 6, 1399–1406 (2011).

13. E. T. Reese, R. G. Siu, H. S. Levinson, The biological degradation of soluble cellulose derivatives and its relationship to the mechanism of cellulose hydrolysis. J. Bacteriol. 59, 485–497 (1950).

14. E. Drula, M. L. Garron, S. Dogan, V. Lombard, B. Henrissat et al., The carbohydrate-active enzyme database: functions and literature. Nucleic Acids Res. 50, D571–D577 (2022).

15. F. Sabbadin, G. R. Hemsworth, L. Ciano, B. Henrissat, P. Dupree et al., An ancient family of lytic polysaccharide monooxygenases with roles in arthropod development and biomass digestion. Nat. Commun. 9, 756 (2018).

16. S. K. Yadav, Archana, R. Singh, P. K. Singh, P. G. Vasudev, Insecticidal fern protein Tma12 is possibly a lytic polysaccharide monooxygenase. Planta 249, 1987–1996 (2019).

17. E. Chiu, M. Hijnen, R. D. Bunker, M. Boudes, C. Rajendran et al., Structural basis for the enhancement of virulence by viral spindles and their in vivo crystallization. Proc. Natl. Acad. Sci. U. S. A. 112, 3973–3978 (2015).

18. T. M. Vandhana, J.-L. Reyre, D. Sushmaa, J.-G. Berrin, B. Bissaro et al., On the expansion of biological functions of lytic polysaccharide monooxygenases. New Phytol. 233, 2380–2396 (2022).

19. E. Wong, G. Vaaje-Kolstad, A. Ghosh, R. Hurtado-Guerrero, P. V. Konarev et al., The Vibrio cholerae colonization factor GbpA possesses a modular structure that governs binding to different host surfaces. PLoS Pathog. 8, e1002373 (2012).

20. S. Garcia-Santamarina, C. Probst, R. A. Festa, C. Ding, A. D. Smith et al., A lytic polysaccharide monooxygenase-like protein functions in fungal copper import and meningitis. Nat. Chem. Biol. 16, 337–344 (2020).

21. Y. Li, X. Y. Liu, M. X. Liu, Y. Wang, Y. B. Zou et al., Magnaporthe oryzae auxiliary activity protein MoAA91 functions as chitin-binding protein to induce appressorium formation on artificial inductive surfaces and suppress plant immunity. mBio. 11, e03304–03319 (2020).

22. F. Askarian, S. Uchiyama, H. Masson, H. V. Sørensen, O. Golten et al., The lytic polysaccharide monooxygenase CbpD promotes Pseudomonas aeruginosa virulence in systemic infection. Nat. Commun. 12, 1230 (2021).

23. F. Sabbadin, B. Henrissat, N. C. Bruce, S. J. McQueen-Mason, Lytic polysaccharide monooxygenases as chitin-specific virulence factors in crayfish plague. Biomolecules 11 (2021).

24. F. Sabbadin, S. Urresti, B. Henrissat, A. O. Avrova, L. R. J. Welsh et al., Secreted pectin monooxygenases drive plant infection by pathogenic oomycetes. Science 373, 774–779 (2021).

25. T. E. Takasuka, A. J. Book, G. R. Lewin, C. R. Currie, B. G. Fox, Aerobic deconstruction of cellulosic biomass by an insect-associated Streptomyces. Sci. Rep. 3, 1030 (2013).

26. C. A. Fowler, F. Sabbadin, L. Ciano, G. R. Hemsworth, L. Elias et al., Discovery, activity and characterisation of an AA10 lytic polysaccharide oxygenase from the shipworm symbiont Teredinibacter turnerae. Biotechnol. Biofuels 12, 232 (2019).

27. A. P. Gonçalves, J. Heller, E. A. Span, G. Rosenfield, H. P. Do et al., Allorecognition upon fungal cell-cell contact determines social cooperation and impacts the acquisition of multicellularity. Curr. Biol. 29, 3006–3017 (2019).

28. X. B. Zhong, L. Zhang, G. P. van Wezel, E. Vijgenboom, D. Claessen, Role for a lytic polysaccharide monooxygenase in cell wall remodeling in Streptomyces coelicolor. mBio. 13, e0045622 (2022).

29. B. Bissaro, Å. K. Røhr, G. Müller, P. Chylenski, M. Skaugen et al., Oxidative cleavage of polysaccharides by monocopper enzymes depends on H_2_O_2_. Nat. Chem. Biol. 13, 1123–1128 (2017).

30. E. D. Hedegård, U. Ryde, Molecular mechanism of lytic polysaccharide monooxygenases. Chem. Sci. 9, 3866–3880 (2018).

31. B. Bissaro, B. Streit, I. Isaksen, V. G. H. Eijsink, G. T. Beckham et al., Molecular mechanism of the chitinolytic peroxygenase reaction. Proc. Natl. Acad. Sci. U. S. A. 117, 1504–1513 (2020).

32. S. M. Jones, W. J. Transue, K. K. Meier, B. Kelemen, E. I. Solomon, Kinetic analysis of amino acid radicals formed in H_2_O_2_-driven Cu(I) LPMO reoxidation implicates dominant homolytic reactivity. Proc. Natl. Acad. Sci. U. S. A. 117, 11916–11922 (2020).

33. B. Wang, Z. Wang, G. J. Davies, P. H. Walton, C. Rovira, Activation of O2 and H_2_O_2_ by lytic polysaccharide monooxygenases. ACS Catal. 10, 12760–12769 (2020).

34. S. Kuusk, V. G. H. Eijsink, P. Väljamäe, The “life-span” of lytic polysaccharide monooxygenases (LPMOs) correlates to the number of turnovers in the reductant peroxidase reaction. J. Biol. Chem. 299, 105094 (2023).

35. M. Torbjörnsson, M. M. Hagemann, U. Ryde, E. D. Hedegård, Histidine oxidation in lytic polysaccharide monooxygenase. J. Biol. Inorg. Chem. 28, 317–328 (2023).

36. H. Østby, T. R. Tuveng, A. A. Stepnov, G. Vaaje-Kolstad, Z. Forsberg et al., Impact of copper saturation on lytic polysaccharide monooxygenase performance. ACS Sustain. Chem. Eng. 11, 15566–15576 (2023).

37. A. A. Stepnov, V. G. H. Eijsink, Z. Forsberg, Enhanced in situ H_2_O_2_ production explains synergy between an LPMO with a cellulose-binding domain and a single-domain LPMO. Sci. Rep. 12, 6129 (2022).

38. D.M. Petrović, B. Bissaro, P. Chylenski, M. Skaugen, M. Sørlie et al., Methylation of the N-terminal histidine protects a lytic polysaccharide monooxygenase from auto-oxidative inactivation. Protein Sci. 27, 1636–1650 (2018).

39. H. B. Gray, J. R. Winkler, Functional and protective hole hopping in metalloenzymes. Chem. Sci. 12, 13988–14003 (2021).

40. J. Zhao, Y. Zhuo, D. E. Diaz, M. Shanmugam, A. J. Telfer et al., Mapping the initial stages of a protective pathway that enhances catalytic turnover by a lytic polysaccharide monooxygenase. J. Am. Chem. Soc. 145, 20672–20682 (2023).

41. I. Ayuso-Fernández, T. Z. Emrich-Mills, J. Haak, O. Golten, K. R. Hall et al., Mutational dissection of a hole hopping route in a lytic polysaccharide monooxygenase (LPMO). Nat. Commun. 15, 3975 (2024).

42. D. N. Bolam, A. Ciruela, S. McQueen-Mason, P. Simpson, M. P. Williamson et al., Pseudomonas cellulose-binding domains mediate their effects by increasing enzyme substrate proximity. Biochem. J. 331, 775–781 (1998).

43. A. B. Boraston, D. N. Bolam, H. J. Gilbert, G. J. Davies, Carbohydrate-binding modules: fine-tuning polysaccharide recognition. Biochem. J. 382, 769–781 (2004).

44. D. Guillen, S. Sanchez, R. Rodriguez-Sanoja, Carbohydrate-binding domains: multiplicity of biological roles. Appl. Microbiol. Biotechnol. 85, 1241–1249 (2010).

45. A. Várnai, M. Siika-Aho, L. Viikari, Carbohydrate-binding modules (CBMs) revisited: reduced amount of water counterbalances the need for CBMs. Biotechnol. Biofuels 6, 30 (2013).

46. G. Courtade, Z. Forsberg, E. B. Heggset, V. G. H. Eijsink, F. L. Aachmann, The carbohydrate-binding module and linker of a modular lytic polysaccharide monooxygenase promote localized cellulose oxidation. J. Biol. Chem. 293, 13006–13015 (2018).

47. A. Chalak, A. Villares, C. Moreau, M. Haon, S. Grisel et al., Influence of the carbohydrate-binding module on the activity of a fungal AA9 lytic polysaccharide monooxygenase on cellulosic substrates. Biotechnol. Biofuels 12, 206 (2019).

48. P. C. Sun, S. V. Valenzuela, P. Chunkrua, F. I. J. Pastor, C. V. F. P. Laurent et al., Oxidized product profiles of AA9 lytic polysaccharide monooxygenases depend on the type of cellulose. ACS Sustain. Chem. Eng. 9, 14124–14133 (2021).

49. Z. Forsberg, G. Courtade, On the impact of carbohydrate-binding modules (CBMs) in lytic polysaccharide monooxygenases (LPMOs). Essays Biochem. 67, 561–574 (2023).

50. W. Gao, T. Li, H. Zhou, J. Ju, H. Yin, Carbohydrate-binding modules enhance H_2_O_2_ tolerance by promoting lytic polysaccharide monooxygenase active site H_2_O_2_ consumption. J. Biol. Chem. 300, 105573 (2023).

51. J. Abramson, J. Adler, J. Dunger, R. Evans, T. Green et al., Addendum: Accurate structure prediction of biomolecular interactions with AlphaFold 3. Nature 636, E4 (2024).

52. Z. Forsberg, A. K. Mackenzie, M. Sørlie, Å. K. Røhr, R. Helland et al., Structural and functional characterization of a conserved pair of bacterial cellulose-oxidizing lytic polysaccharide monooxygenases. Proc. Natl. Acad. Sci. U. S. A. 111, 8446–8451 (2014).

53. B. Bissaro, E. Kommedal, Å. K. Røhr, V. G. H. Eijsink, Controlled depolymerization of cellulose by light-driven lytic polysaccharide oxygenases. Nat. Commun. 11, 890 (2020).

54. Z. Forsberg, A. A. Stepnov, G. Tesei, Y. Wang, E. Buchinger et al., The effect of linker conformation on performance and stability of a two-domain lytic polysaccharide monooxygenase. J. Biol. Chem. 299, 105262 (2023).

55. A. Carletti, F. Csarman, M. Sola, G. Battistuzzi, R. Ludwig et al., Kinetic and substrate specificity determination of bacterial LPMOs. ACS Catal. 14, 14586–14594 (2024).

56. K. Peariso, D. L. Huffman, J. E. Penner-Hahn, T. V. O’Halloran, The PcoC copper resistance protein coordinates Cu(I) via novel S-methionine interactions. J. Am. Chem. Soc. 125, 342–343 (2003).

57. J. T. Rubino, P. Riggs-Gelasco, K. J. Franz, Methionine motifs of copper transport proteins provide general and flexible thioether-only binding sites for Cu(I) and Ag(I). J. Biol. Inorg. Chem. 15, 1033–1049 (2010).

58. Y. Xue, A. V. Davis, G. Balakrishnan, J. P. Stasser, B. M. Staehlin et al., Cu(I) recognition via cation-pi and methionine interactions in CusF. Nat. Chem. Biol. 4, 107–109 (2008).

59. L. Zhang, M. Koay, M. J. Maher, Z. Xiao, A. G. Wedd, Intermolecular transfer of copper ions from the CopC protein of Pseudomonas syringae. Crystal structures of fully loaded Cu(I)Cu(II) forms. J. Am. Chem. Soc. 128, 5834–5850 (2006).

60. Z. Forsberg, B. Bissaro, J. Gullesen, B. Dalhus, G. Vaaje-Kolstad et al., Structural determinants of bacterial lytic polysaccharide monooxygenase functionality. J. Biol. Chem. 293, 1397–1412 (2018).

61. K. R. Hall, M. Mollatt, Z. Forsberg, O. Golten, L. Schwaiger et al., Impact of the copper second coordination sphere on catalytic performance and substrate specificity of a bacterial lytic polysaccharide monooxygenase. ACS Omega 9, 23040–23052 (2024).

62. A. J. Book, R. M. Yennamalli, T. E. Takasuka, C. R. Currie, G. N. Phillips et al., Evolution of substrate specificity in bacterial AA10 lytic polysaccharide monooxygenases. Biotechnol. Biofuels 7, 109 (2014).

63. A. K. Votvik, Å. K. Røhr, B. Bissaro, A. A. Stepnov, M. Sørlie et al., Structural and functional characterization of the catalytic domain of a cell-wall anchored bacterial lytic polysaccharide monooxygenase from Streptomyces coelicolor. Sci. Rep. 13, 5345 (2023).

64. J. Jumper, R. Evans, A. Pritzel, T. Green, M. Figurnov et al., Highly accurate protein structure prediction with AlphaFold. Nature 596, 583–589 (2021).

65. T. R. Tuveng, M. S. Jensen, L. Fredriksen, G. Vaaje-Kolstad, V. G. H. Eijsink et al., A thermostable bacterial lytic polysaccharide monooxygenase with high operational stability in a wide temperature range. Biotechnol. Biofuels 13, 194 (2020).

66. A. A. Stepnov, Z. Forsberg, M. Sørlie, G. S. Nguyen, A. Wentzel et al., Unraveling the roles of the reductant and free copper ions in LPMO kinetics. Biotechnol. Biofuels 14, 28 (2021).

67. V. A. Rapisarda, S. I. Volentini, R. N. Farias, E. M. Massa, Quenching of bathocuproine disulfonate fluorescence by Cu(I) as a basis for copper quantification. Anal. Biochem. 307, 105–109 (2002).

68. S. Ogawa, R. Ichiki, M. Abo, E. Yoshimura, Revision of analytical conditions for determining ligand molecules specific to soft metal ions using dequenching of copper(I)-bathocuproine disulfonate as a detection system. Anal. Chem. 81, 9199–9200 (2009).

69. R. Kittl, D. Kracher, D. Burgstaller, D. Haltrich, R. Ludwig, Production of four Neurospora crassa lytic polysaccharide monooxygenases in Pichia pastoris monitored by a fluorimetric assay. Biotechnol. Biofuels 5, 79 (2012).

70. M. M. Hagemann, E. K. Wieduwilt, U. Ryde, E. D. Hedegård, Investigating the substrate oxidation mechanism in lytic polysaccharide monooxygenase: H_2_O_2_-versus O2-activation. Inorg. Chem. 63, 21929–21940 (2024).

71. E. K. Wieduwilt, M. M. Hagemann, U. Ryde, E. D. Hedegård, The mechanism behind the oxidase activity of cellulose-active AA10 lytic polysaccharide monooxygenases. Inorg. Chem. Front. 10.1039/d5qi00796h (2025).

72. M. Blum, A. Andreeva, L. C. Florentino, S. R. Chuguransky, T. Grego et al., InterPro: the protein sequence classification resource in 2025. Nucleic Acids Res. 53, D444–D456 (2025).

73. L. Fu, B. Niu, Z. Zhu, S. Wu, W. Li, CD-HIT: accelerated for clustering the next-generation sequencing data. Bioinformatics 28, 3150–3152 (2012).

74. M. Varadi, S. Anyango, M. Deshpande, S. Nair, C. Natassia et al., AlphaFold Protein Structure Database: massively expanding the structural coverage of protein-sequence space with high-accuracy models. Nucleic Acids Res. 50, D439–D444 (2022).

75. M. Mirdita, K. Schutze, Y. Moriwaki, L. Heo, S. Ovchinnikov et al., ColabFold: making protein folding accessible to all. Nat. Methods 19, 679–682 (2022).

76. J. Zheng, Q. Ge, Y. Yan, X. Zhang, L. Huang et al., dbCAN3: automated carbohydrate-active enzyme and substrate annotation. Nucleic Acids Res. 51, W115–W121 (2023).

77. K. Katoh, K. Misawa, K. Kuma, T. Miyata, MAFFT: a novel method for rapid multiple sequence alignment based on fast Fourier transform. Nucleic Acids Res. 30, 3059–3066 (2002).

78. M. N. Price, P. S. Dehal, A. P. Arkin, FastTree: computing large minimum evolution trees with profiles instead of a distance matrix. Mol. Biol. Evol. 26, 1641–1650 (2009).

79. C. W. Simmons, A. P. Reddy, P. D’Haeseleer, J. Khudyakov, K. Billis et al., Metatranscriptomic analysis of lignocellulolytic microbial communities involved in high-solids decomposition of rice straw. Biotechnol. Biofuels 7, 495 (2014).

80. C. Manoil, J. Beckwith, A genetic approach to analyzing membrane protein topology. Science 233, 1403–1408 (1986).

81. D. C. Irwin, M. Spezio, L. P. Walker, D. B. Wilson, Activity studies of eight purified cellulases: specificity, synergism, and binding domain effects. Biotechnol. Bioeng. 42, 1002–1013 (1993).

82. B. Westereng, J. W. Agger, S. J. Horn, G. Vaaje-Kolstad, F. L. Aachmann et al., Efficient separation of oxidized cello-oligosaccharides generated by cellulose degrading lytic polysaccharide monooxygenases. J. Chromatogr. A. 1271, 144–152 (2013).

83. B. Bissaro, Z. Forsberg, Y. Ni, F. Hollmann, G. Vaaje-Kolstad et al., Fueling biomass-degrading oxidative enzymes by light-driven water oxidation. Green Chem. 18, 5357–5366 (2016).

