## Supplementary Table S1 and Figures S1 to S12 for "Discovery and characterization of a copper-binding carbohydrate-binding module (CBM) regulating the activity of lytic polysaccharide monooxygenases"

**This PDF file includes:**

Table S1
Figures S1 to S12

**Table S1. Copper content for variants of *Sc*LPMO10C and mgLPMO10 determined by ICP-MS.** After copper saturation of all AA10-containing variants, the copper content of 2 µM protein was analyzed using ICP-MS. CBM2s were not preloaded with copper before the analysis. The copper content of the two mutated CBMs (i.e., *Sc*CBM2$^{AAA}$ and mgCBM2$^{MMH}$) was not determined (n.d.), as the same mutations in the full-length enzyme did not affect the copper content of these enzymes. Standard deviations for duplicate sample preparations are shown (n=2).

| Enzyme/protein | Copper content (µM) | Ratio (Cu:protein) |
|---|---|---|
| *Sc*LPMO10C$^{(MMH)}$ | 1.55 ± 0.21 | 0.775:1 |
| *Sc*AA10 | 1.50 ± 0.00 | 0.750:1 |
| *Sc*CBM2$^{(MMH)}$ | 0.06 ± 0.03 | 0.032:1 |
| *Sc*AA10-mgCBM2$^{(ART)}$ | 1.60 ± 0.14 | 0.800:1 |
| *Sc*LPMO10C$^{AAA}$ | 1.75 ± 0.07 | 0.875:1 |
| *Sc*CBM2$^{AAA}$ | n.d. | n.d. |
| mgLPMO10$^{(ART)}$ | 1.20 ± 0.00 | 0.600:1 |
| mgAA10 | 1.25 ± 0.07 | 0.625:1 |
| mgCBM2$^{(ART)}$ | 0.03 ± 0.00 | 0.015:1 |
| mgAA10-ScCBM2$^{(MMH)}$ | 1.30 ± 0.00 | 0.650:1 |
| mgLPMO10$^{MMH}$ | 1.20 ± 0.14 | 0.600:1 |
| mgCBM2$^{MMH}$ | n.d. | n.d. |

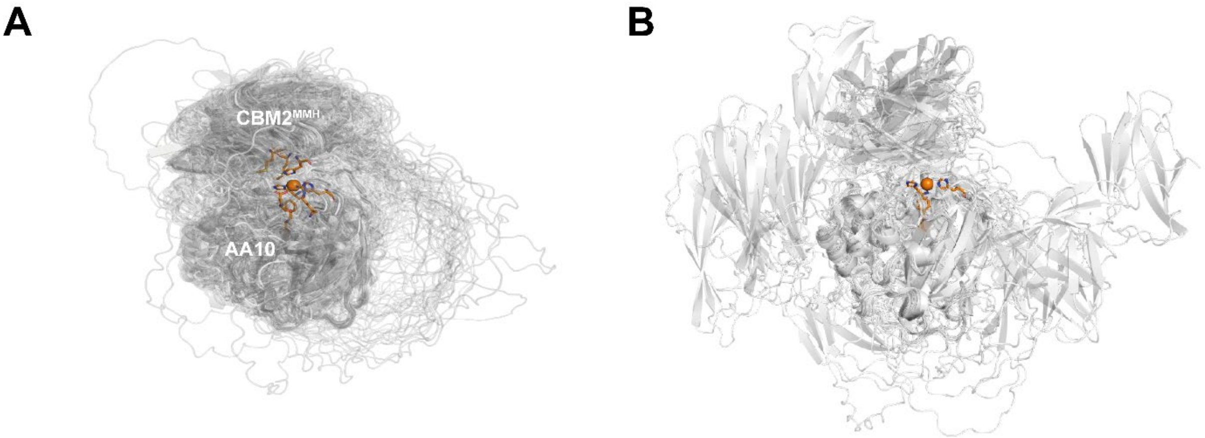


**Figure S1. Structural alignment of CBM2 containing LPMOs.** Panel A shows a superposition of AlphaFold models for the 130 MMH-containing two-domain LPMOs identified in the phylogenetic analysis shown in Figure 2B of the main manuscript (i.e., all LPMOs labeled green). Panel B shows the predicted structures of 28 sequences of CBM2 containing AA10 LPMOs lacking the MMH motif, which display more variable orientations of the CBM2 relative to the catalytic domain. The models were generated using AlphaFold2 [1] and were aligned and visualized in PyMOL.

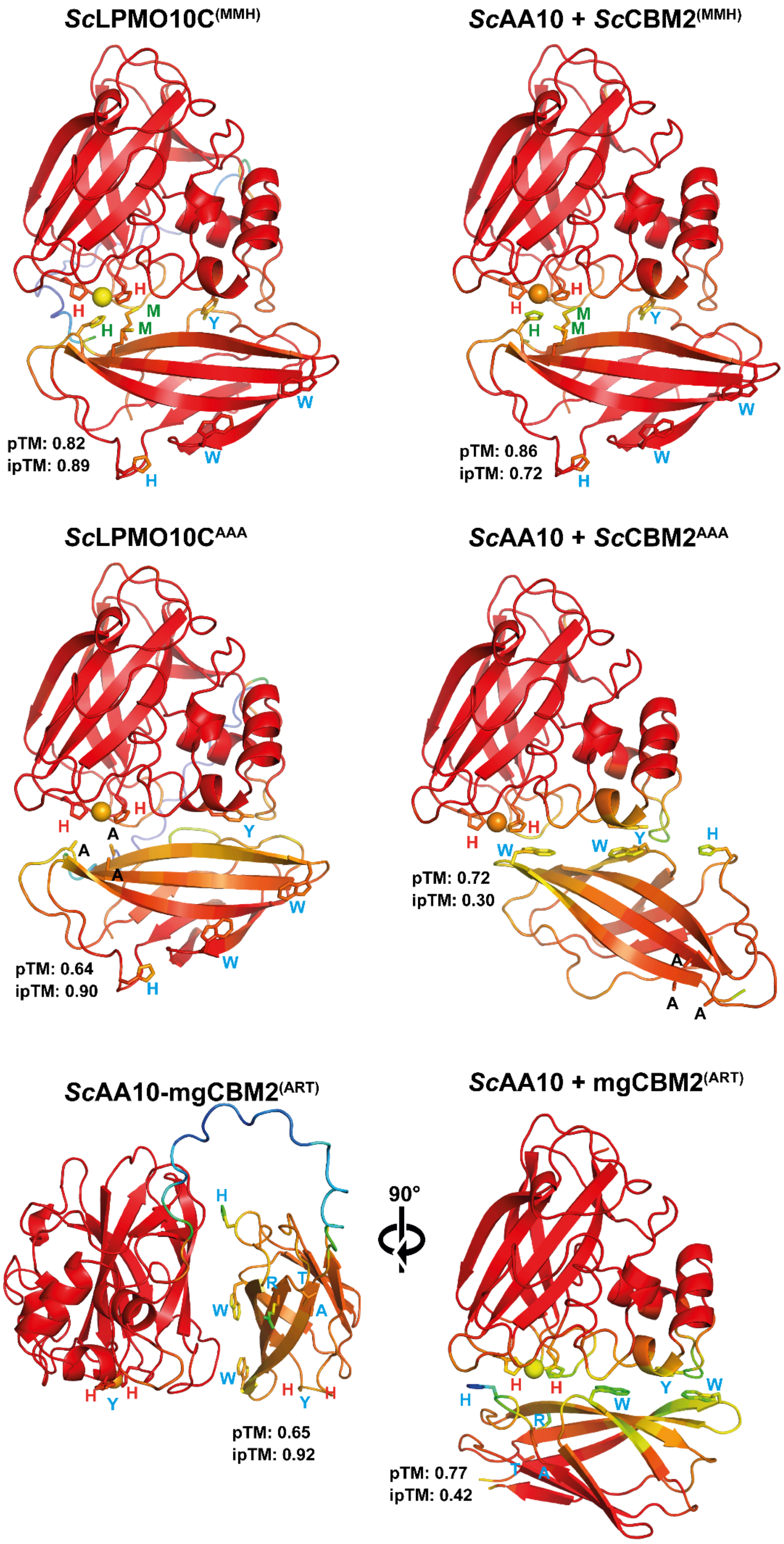
ScLPMO10C(MMH)
pTM: 0.82
ipTM: 0.89
ScAA10 + ScCBM2(MMH)
pTM: 0.86
ipTM: 0.72
ScLPMO10CAAA
pTM: 0.64
ipTM: 0.90
ScAA10 + ScCBM2AAA
pTM: 0.72
ipTM: 0.30
ScAA10-mgCBM2(ART)
pTM: 0.65
ipTM: 0.92
90°
ScAA10 + mgCBM2(ART)
pTM: 0.77
ipTM: 0.42

**Figure S2. Structural representation of the interaction between *Sc*AA10 and different CBM2 variants based on AlphaFold3 models.** Each pair of structures shows the predicted intramolecular (linker present; left) and intermolecular (linker not present, right) interaction between the CD and the CBM2. Residues are colored according to predicted Local Distance Difference Test (pLDDT) scores, ranging from blue (low confidence) to red (high confidence). The predicted Template Modeling (pTM) score reflects confidence in the global arrangement of domains, with values near 1.0 indicating high reliability. For complexes, the interface predicted Template Modeling (ipTM) score assesses confidence in domain interactions. Note that only three of these structures, *Sc*LPMO10C$^{(MMH)}$, *Sc*AA10 + *Sc*CBM2$^{(MMH)}$ and *Sc*LPMO10C$^{AAA}$, show an interaction between the catalytic copper site and the wildtype or mutated MMH site. In two other cases, complex formation is also observed, but with much lower reliability and not involving the MMH site. When substituting the CBM2 domain (as in *Sc*AA10-mgCBM2$^{(ART)}$), the interaction does not occur. Residues for which side chains are shown are those that are discussed in the main text and are labeled using the single-letter amino acid code.

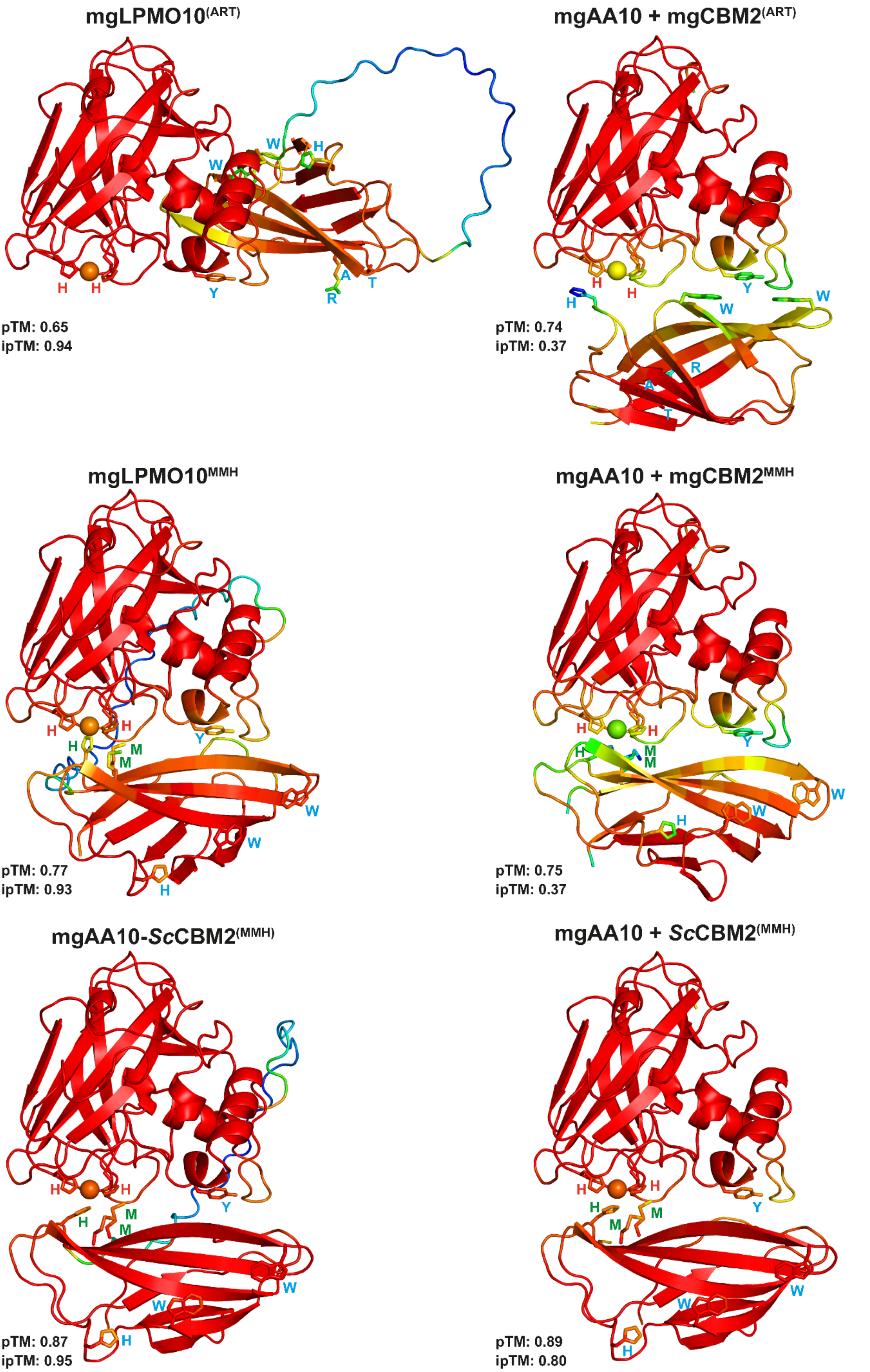
mgLPMO10(ART)
W
H
W
H
H
Y
A
T
R
pTM: 0.65
ipTM: 0.94
mgAA10 + mgCBM2(ART)
H
H
Y
H
W
W
R
A
T
pTM: 0.74
ipTM: 0.37
mgLPMO10MMH
H
H
H
M
M
Y
W
W
H
pTM: 0.77
ipTM: 0.93
mgAA10 + mgCBM2MMH
H
H
H
M
M
Y
W
W
H
pTM: 0.75
ipTM: 0.37
mgAA10-ScCBM2(MMH)
H
H
H
M
M
Y
W
W
H
pTM: 0.87
ipTM: 0.95
mgAA10 + ScCBM2(MMH)
H
H
H
M
M
Y
W
W
H
pTM: 0.89
ipTM: 0.80

**Figure S3. Structural representation of the interaction between mgAA10 and CBM2 variants based on AlphaFold3 models**. Each pair of structures shows the predicted intramolecular (linker present; left) and intermolecular (linker not present, right) interaction between the CD and the CBM2. Residue confidence is shown by pLDDT scores (blue: low, red: high). The predicted pTM score indicates global domain arrangement reliability, and ipTM scores reflect interaction confidence (see legend to Fig. S2 for more details). Residues for which side chains are shown are those that are discussed in the main text and are labeled using the single-letter amino acid code. Note that only structures in which the MMH motif is present show the interaction between this motif and the catalytic copper site. It is also worth noting that the presence of the linker improves the ipTM.

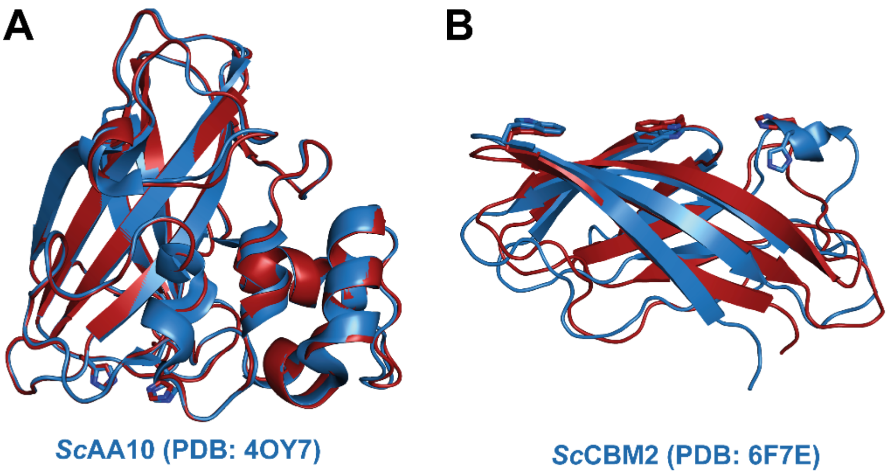


**Figure S4. Structural comparison of experimentally determined and predicted structures.** The figure compares the crystal structures of *Sc*AA10 (A) (PDB: 4OY7 [2]) and the NMR structure of *Sc*CBM2 (B) (PDB: 6F7E [3]) with AlphaFold 3 structure predictions. Experimentally determined structures are shown in blue, and AlphaFold 3-predicted models in red. The structural alignment resulted in a low RMSD of 0.26 Å for the AA10 crystal structure, indicating high accuracy in the predicted model. For the CBM2 domain, which was solved using NMR spectroscopy, the RMSD is higher (1.56 Å) due to structural variability across the 20 conformers in the NMR ensemble. This is expected, as NMR ensembles capture conformational flexibility, leading to higher RMSD values when compared to a single predicted model.

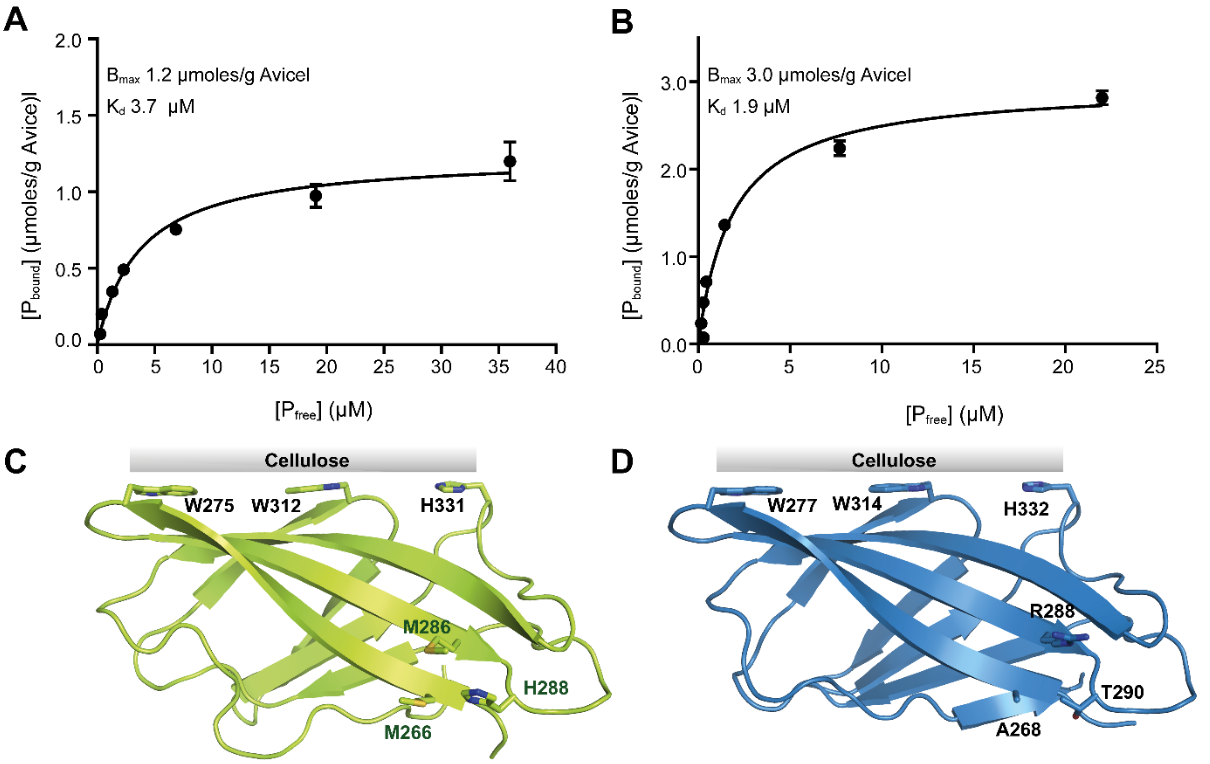


**Figure S5. Binding of *Sc*CBM2$^{(MMH)}$ and mgCBM2$^{(ART)}$ to Avicel.** The plots show binding data for *Sc*CBM2$^{(MMH)}$ (A) and mgCBM2$^{(ART)}$ (B) incubated with Avicel for 60 min. The experiments were carried out at 22 °C using 10 g/L Avicel in 50 mM sodium phosphate buffer (pH 6.0). $P_{bound}$ corresponds to bound protein (µmoles/g substrate), and $P_{free}$ corresponds to non-bound protein (µM). The error bars show ± SD (n = 3). Panels C and D show the substrate binding residues in *Sc*CBM2 (C; Trp275, Trp312 & His331) and mgCBM2 (D; Trp277, Trp314 & His332) and the relative location to the predicted copper binding site in *Sc*CBM2 (Met266, Met286 & His288) and equivalent residues in mgCBM2 (Ala268, Arg288 & Thr290).

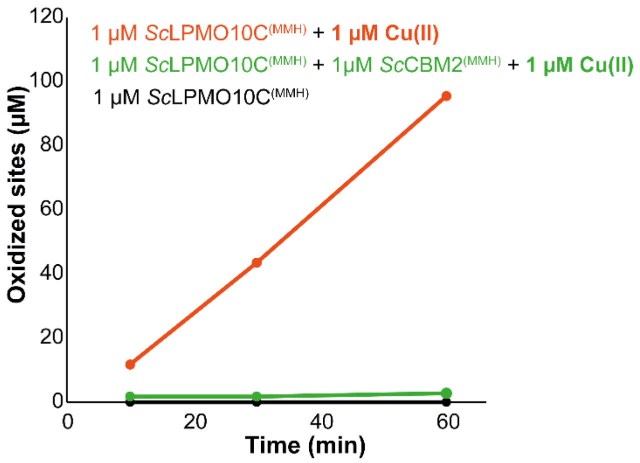


**Figure S6. Initial rate of the LPMO reaction with cellulose measured in the presence and absence of excess copper.** Reactions containing 1 µM copper saturated *Sc*LPMO10C$^{(MMH)}$ were incubated with 10 g/L Avicel at 40°C in 50 mM sodium phosphate buffer (pH 6.0), with 1 mM AscA, for up to 60 min, in the presence or absence of 1 µM Cu(II)$SO_4$, with or without 1 µM *Sc*CBM2$^{(MMH)}$. At various time points, samples were taken, and reactions were stopped by vacuum filtration. The soluble oxidized products were then converted to oxidized dimers and trimers by treatment with the *Thermobifida fusca* endoglucanase Cel6A (*Tf*Cel6A) followed by chromatographic analysis and quantification. "Oxidized sites" represents the sum of oxidized dimers and trimers. The progress curves show that, as expected, the presence of free copper leads to massively increased LPMO activity, because copper promotes abiotic oxidation of ascorbic acid, which will generate $H_2O_2$ [4]. This effect of free copper is not observed in the presence of the copper-binding CBM2.

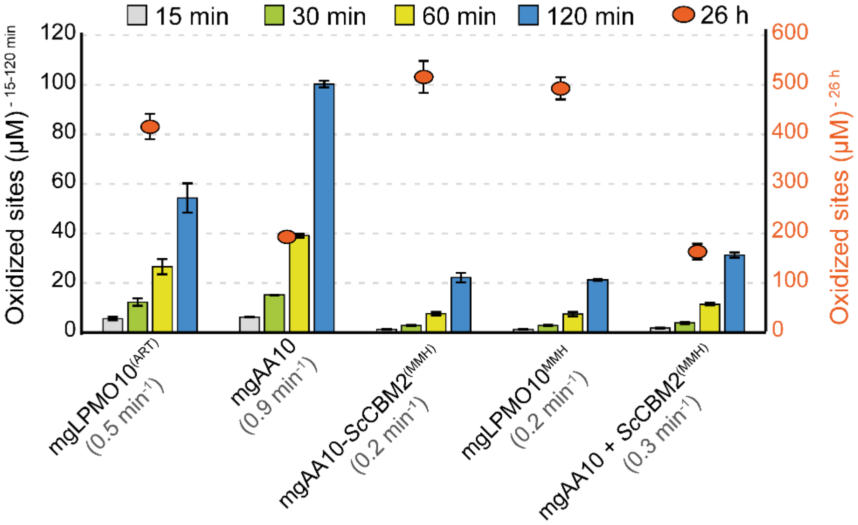


**Figure S7. Progress curves for degradation of Avicel by mgLPMO10 variants and various combinations of individual domains.** Reactions containing 1 µM LPMO were incubated with 10 g/L Avicel at 40°C in 50 mM sodium phosphate buffer (pH 6.0), with 1 mM ascorbic acid, for up to 26 hours. At various time points, samples were taken, and reactions were stopped by vacuum filtration. The soluble oxidized products were then converted to oxidized dimers and trimers by treatment with *Tf*Cel6A, which were quantified to yield oxidized sites (left y-axis for the early time points; right y-axis for the 26 h time point). The initial rates (shown in brackets for each reaction) were estimated from the linear phase of the reaction. Note that under the conditions used here, the LPMO reaction is limited by the rate of *in situ* generation of $H_2O_2$. Error bars represent standard deviations (n = 3).

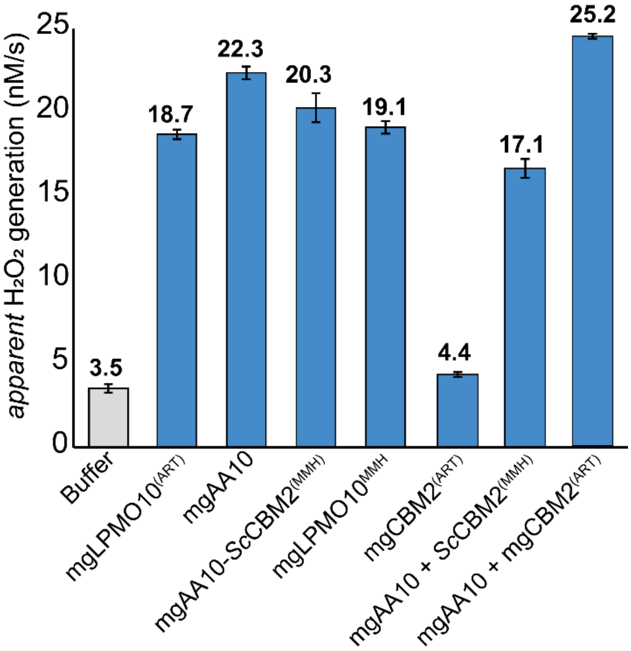


**Figure S8. Oxidase activity of mgLPMO10 variants measured using the Amplex Red/HRP assay.** The bar chart shows the apparent rates of $H_2O_2$ production in various reactions containing various LPMOs or (combinations of) LPMO domains (blue bars). A buffer control is shown for comparison (grey bar). All reactions were performed with 4 µM LPMO, with or without 4 µM CBM2, in 50 mM sodium phosphate buffer (pH 6.0). The reaction mixture contained 1 mM ascorbic acid, 5 U/mL HRP, 100 µM Amplex Red, and 1% (v/v) DMSO. Reaction rates were determined using the linear phase of the reaction (approximately 0–120 min), and error bars represent ± standard deviation (n = 3).

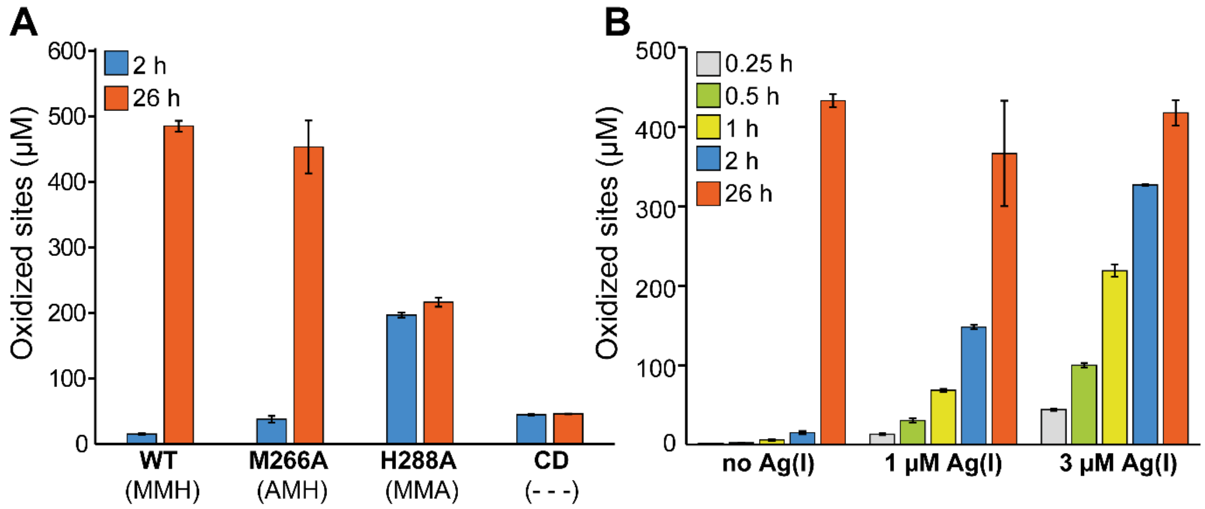


**Figure S9. Effects of modulation of the copper-binding ability of the CBM2 in *Sc*LPMO10C$^{(MMH)}$ on cellulose degradation.** Panel A shows Avicel oxidation by wildtype *Sc*LPMO10C$^{(MMH)}$, two single mutants (M266A and H288A) and the catalytic domain only. The copper-binding motifs in the CBM are indicated in brackets for clarity. Panel B shows the quantification of soluble products over time from Avicel degradation by the wildtype enzyme pre-incubated with 0, 1, or 3 µM Ag(I)$NO_3$ before starting the reaction. All reactions contained 1 µM LPMO and were incubated with 10 g/L Avicel at 40°C in 50 mM sodium phosphate buffer (pH 6.0), with 1 mM ascorbic acid, for up to 26 hours. At various time points, samples were taken, and reactions were stopped by vacuum filtration. The soluble oxidized products were then converted to oxidized dimers and trimers by treatment with *Tf*Cel6A which were quantified to yield oxidized sites. Error bars represent standard deviations (n = 3).

| CBM motif | MMH | MMHH | MMH | MHH | MHMM |
|---|---|---|---|---|---|
| 2nd sphere motif | REF | REF | REF | REF | HQF* |
| ***Relative location CBM2 Cu-site*** | *"N-terminal"* | *"N-terminal and central region"* | *"C-terminal"* | *"central region"* | *"central region"* |
| **Organism** | *Streptomyces coelicolor* A3(2) | *Actinoplanes friuliensis* | *Streptomyces malaysiensis* | *Cellulomonas gelida* | *Glycomyces artemisiae* |
| **Uniprot ID** | Q9RJY2 | U5W274 | QDL70221 | A0A4Y3KSF6 | A0A2T0UIF0 |

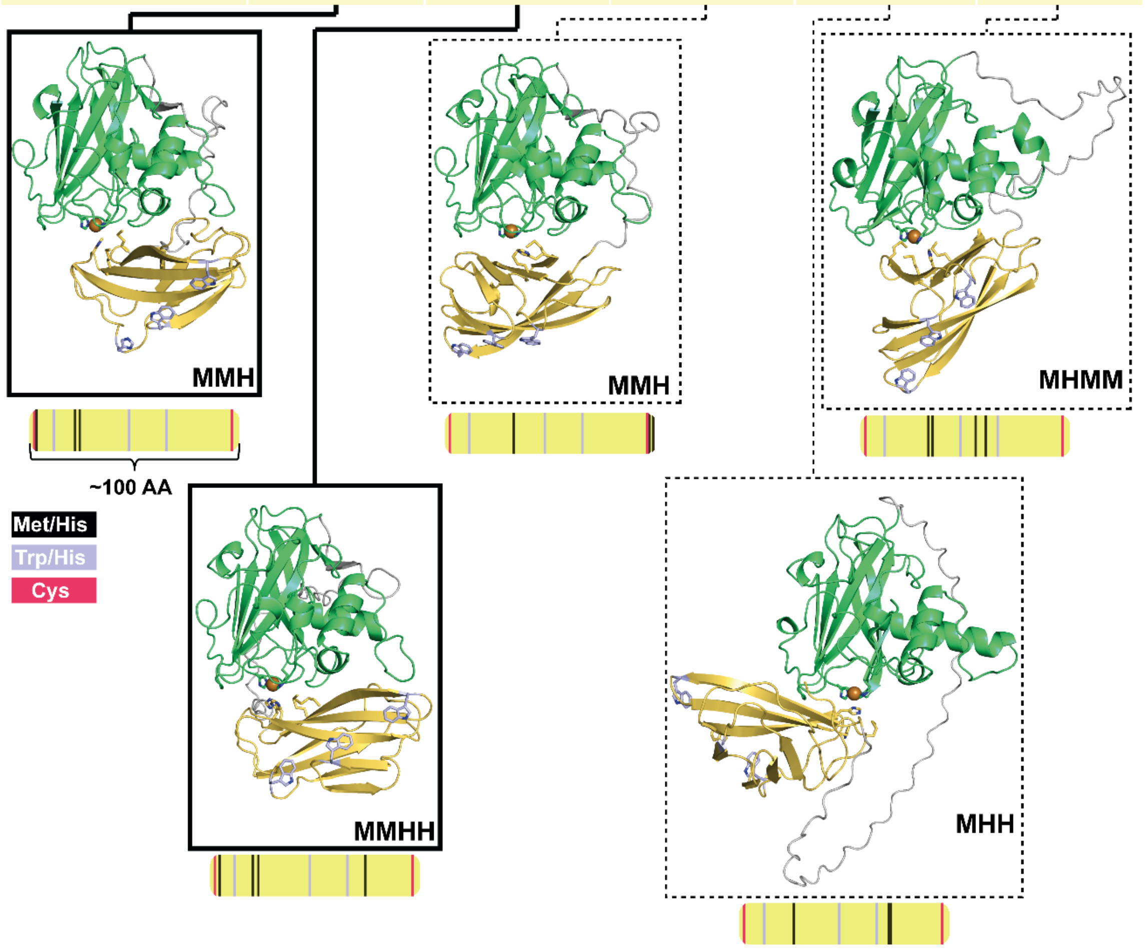


**Figure S10. Putative interactions between LPMO domains (green) and appended copper-binding CBM2 domains (yellow).** The figure illustrates five distinct Met-His (MH)-containing motifs within CBM2 domains that may be involved in copper coordination. The first two motifs, MMH and MMHH (black solid lines), have been experimentally confirmed to bind copper in this study. The three other putative sites (indicated by dashed lines), include a second MMH motif, identified in the *S. malaysiensis* LPMO, with a methionine in the central region of the CBM and a C-terminal Met-His pair. A MHH motif is observed in multiple *Cellulomonas* LPMO sequences, including an LPMO from *C. gelida*, which is shown here. The fifth motif, MHMM, is found in an AA10 that differs from the other AA10s with a copper-binding CBM2 in that it does not contain the REF second sphere motif. This LPMO falls within the approximately 1% of sequences categorized as "other than REF" in Fig. 2A and its second sphere arrangement, HQF (labeled with an asterisk), appears to represent an intermediate between REF-type and HQY-type AA10 LPMOs. Note the varying spatial orientation (i.e., linker length and shape) of the CBM relative to the catalytic domain in the AlphaFold3 models. This variation appears to be influenced by the position of the putative copper-binding methionine and histidine residues on the CBM surface. The yellow bars depict the ~100-residue CBM2 domain, highlighting the relative positions of the fully conserved cellulose-binding residues (purple) and the N- and C-terminal cysteines (pink) that form a disulfide bridge. The putative copper-binding residues are shown in black, and their positions vary across the CBMs, contributing to the observed differences in CBM-CD orientation and interaction.

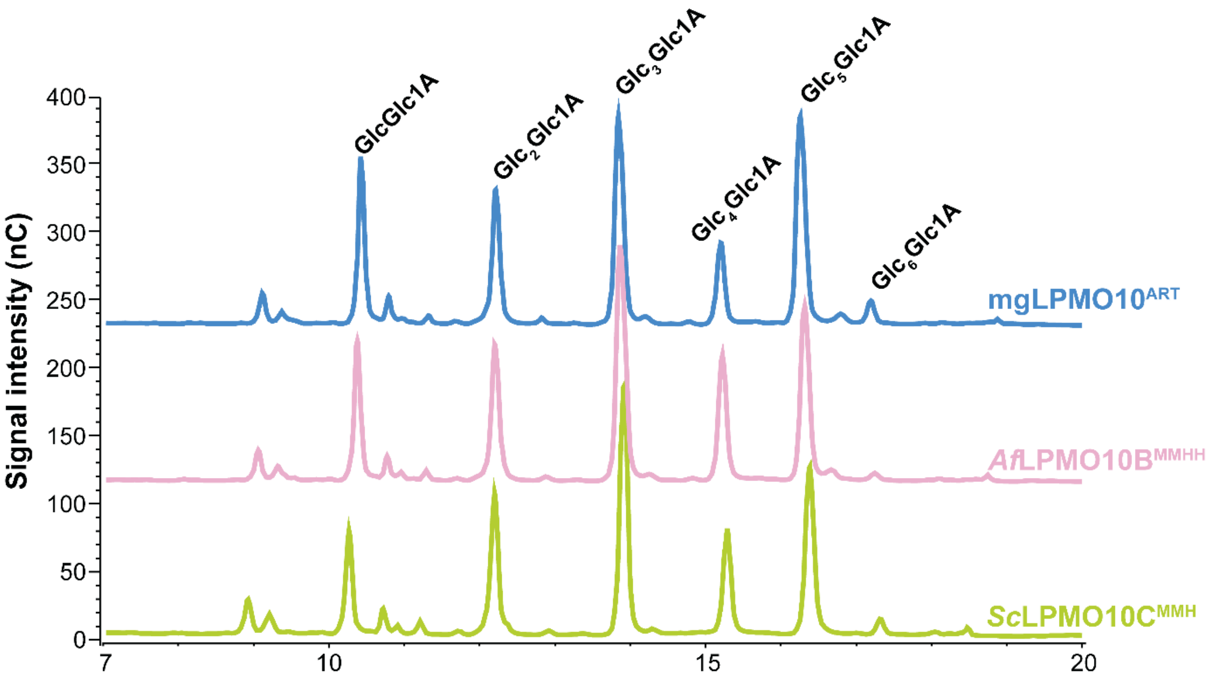


**Figure S11. Product profiles of three wildtype LPMOs.** High-performance anion-exchange chromatography with pulsed amperometric detection (HPAEC-PAD) was used to analyze the oxidized products generated by the three wildtype LPMOs in this study (mgLPMO10$^{(ART)}$, *Af*LPMO10B$^{(MMHH)}$ and *Sc*LPMO10C$^{(MMH)}$). The data confirm that all enzymes exhibit similar activity, that is C1-oxidation of cellulose. Reactions were conducted with 1 µM LPMO and 10 g/L Avicel in 50 mM sodium phosphate buffer (pH 6.0) at 40 °C, with 1 mM AscA as reductant. After 26 hours of incubation, reaction products were analyzed.

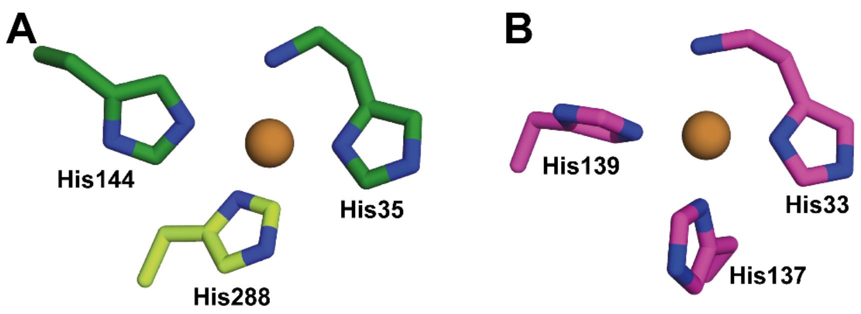


**Figure S12. Structural resemblance between the copper site in the LPMO-CBM2 complex and the Cu(B) site in particulate methane monooxygenases (pMMOs).** The copper site in the LPMO–CBM2 complex (panel A) features an additional histidine ligand (His288 from the CBM2) relative to the canonical LPMO active site (AlphaFold3 predicted structure), forming a coordination environment that closely resembles the Cu(B) site (panel B) found in pMMOs (PDB code 3RGB; [5]). In both systems, the copper ion is coordinated by three histidine residues in a similar spatial arrangement. This resemblance may reflect an analogous adaptation for stabilizing copper in a redox active environment. Structures were aligned based on the copper-binding residues. Note that His35 (A) and His33 (B) are the N-terminal residues of the proteins, interacting with the copper through both the imidazole side chain and the N-terminal amino group.

### References

1. J. Jumper, R. Evans, A. Pritzel, T. Green, M. Figurnov *et al.*, Highly accurate protein structure prediction with AlphaFold. *Nature* **596**, 583-589 (2021).
2. Z. Forsberg, A. K. Mackenzie, M. Sørlie, Å. K. Røhr, R. Helland *et al.*, Structural and functional characterization of a conserved pair of bacterial cellulose-oxidizing lytic polysaccharide monooxygenases. *Proc. Natl. Acad. Sci. U. S. A.* **111**, 8446-8451 (2014).
3. G. Courtade, Z. Forsberg, E. B. Heggset, V. G. H. Eijsink, F. L. Aachmann, The carbohydrate-binding module and linker of a modular lytic polysaccharide monooxygenase promote localized cellulose oxidation. *J. Biol. Chem.* **293**, 13006-13015 (2018).
4. A. A. Stepnov, Z. Forsberg, M. Sørlie, G. S. Nguyen, A. Wentzel *et al.*, Unraveling the roles of the reductant and free copper ions in LPMO kinetics. *Biotechnol. Biofuels* **14**, 28 (2021).
5. S. M. Smith, S. Rawat, J. Telser, B. M. Hoffman, T. L. Stemmler *et al.*, Crystal structure and characterization of particulate methane monooxygenase from *Methylocystis* species strain M. *Biochemistry-Us* **50**, 10231-10240 (2011).
